# ITPK1-Dependent Inositol Polyphosphates Regulate Auxin Responses in *Arabidopsis thaliana*

**DOI:** 10.1101/2020.04.23.058487

**Authors:** Nargis Parvin Laha, Yashika Walia Dhir, Ricardo F.H. Giehl, Eva Maria Schäfer, Philipp Gaugler, Zhaleh Haghighat Shishavan, Hitika Gulabani, Haibin Mao, Ning Zheng, Nicolaus von Wirén, Henning J. Jessen, Adolfo Saiardi, Saikat Bhattacharjee, Debabrata Laha, Gabriel Schaaf

## Abstract

The combinatorial phosphorylation of *myo-inositol* results in the generation of different inositol phosphates (InsP), of which phytic acid (InsP_6_) is the most abundant species in eukaryotes. InsP_6_ is also the precursor of higher phosphorylated forms called inositol pyrophosphates (PP-InsPs), such as InsP_7_ and InsP_8_, which are characterized by a diphosphate moiety and are also ubiquitously found in eukaryotic cells. While PP-InsPs regulate various cellular processes in animals and yeast, their biosynthesis and functions in plants has remained largely elusive because plant genomes do not encode canonical InsP_6_ kinases. Recently, it was shown that Arabidopsis ITPK1 catalyzes the phosphorylation of InsP_6_ to the natural 5-InsP_7_ isomer *in vitro*. Here, we demonstrate that Arabidopsis ITPK1 contributes to the synthesis of InsP_7_ *in planta*. We further find a critical role of ITPK1 in auxin-related processes including primary root elongation, leaf venation, thermomorphogenic and gravitropic responses, and sensitivity towards exogenously applied auxin. Notably, 5-InsP_7_ binds to recombinant auxin receptor complex, consisting of the F-Box protein TIR1, ASK1 and the transcriptional repressor IAA7, with high affinity. Furthermore, a specific increase in 5-InsP_7_ in a heterologous yeast expression system results in elevated interaction of the TIR1 homologs AFB1 and AFB2 with various AUX/IAA-type transcriptional repressors. We also identified a physical interaction between ITPK1 and TIR1, suggesting a dedicated channeling of an activating factor, such as 5-InsP_7_, to the auxin receptor complex. Our findings expand the mechanistic understanding of auxin perception and lay the biochemical and genetic basis to uncover physiological processes regulated by 5-InsP_7_.

## INTRODUCTION

The phytohormone auxin orchestrates a plethora of growth and developmental processes, including embryogenesis, root development and gravitropism (Teale et al., 2006; Lavenus et al., 2013; Salehin et al., 2015; Weijers and Wagner, 2016). Its distribution within plant tissues creates various organized patterns, such as leaf venation (Scarpella et al., 2006), phyllotactic patterns (Reinhardt et al., 2003; Jönsson et al., 2006; Hartmann et al., 2019) and xylem differentiation (Fukuda and Komamine, 1980; Bishopp et al., 2011; Smetana et al., 2019). Auxin perception is mediated by TRANSPORT INHIBITOR RESPONSE1 (TIR1) and AUXIN-SIGNALING F-BOX proteins (AFB1-5), which induce SCF ubiquitin-ligase-catalyzed degradation of Aux/IAA transcriptional repressors to activate AUXIN RESPONSE FACTOR (ARF) transcription factors (Gray et al., 2001; Dharmasiri et al., 2005; Kepinski and Leyser, 2005; Prigge et al., 2020). Unexpectedly, in a crystal structure of the auxin receptor complex consisting of insect-purified ASK1-TIR1 and an IAA7 degron peptide, insect-derived InsP_6_ occupied the core of the leucine-rich-repeat (LRR) domain of TIR1 (Tan et al., 2007). While the functional importance of InsP_6_ in auxin perception remains elusive, this molecule serves as a major phosphate store in seeds and as precursor of InsP_7_ and InsP_8_, in which the *myo*-inositol ring contains one or more energy-rich diphosphate moieties.

PP-InsPs regulate a wide range of important biological functions, such as vesicular trafficking, ribosome biogenesis, immune response, DNA repair, telomere length maintenance, phosphate homeostasis, spermiogenesis, insulin signaling and cellular energy homeostasis in yeast and mammals (Wilson et al., 2013; Thota et al., 2015; Wild et al., 2016; Shears, 2017; Wilson et al., 2019). PP-InsPs were also identified in different plant species (Brearley and Hanke, 1996; Flores and Smart, 2000; Dorsch et al., 2003; Desai et al., 2014; Laha et al., 2015). They have been shown to represent potential targets of bacterial type III effector InsP hydrolytic enzymes in pepper and tomato (Blüher et al., 2017), to act as regulators of TOR signaling in Chlamydomonas (Couso et al., 2016), as well as to represent critical co-ligands of the CORONATINE INSENSITIVE 1/JASMONATE ZIM DOMAIN (COI1-JAZ) jasmonate receptor complex (Laha et al., 2015; Laha et al., 2016). PP-InsPs were also shown to regulate phosphate sensing in Arabidopsis (Wild et al., 2016; Dong et al., 2019; Zhu et al., 2019) by promoting the physical interaction between PHOSPHATE STARVATION RESPONSE (PHR) transcription factors with stand-alone SYG1/Pho81/XPR1 (SPX) proteins (Wild et al., 2016; Dong et al., 2019). Upon interaction with PHRs, SPX proteins inhibit PHR-dependent activation of phosphate deficiency-induced genes, thereby preventing phosphate over-accumulation under conditions of high phosphate availability (Lv et al., 2014; Puga et al., 2014; Wang et al., 2014). Because of the low binding affinities of phosphate ions to the PHR1/SPX1 complex, a direct role of phosphate itself in regulating this interaction is rather unlikely (Lv et al., 2014; Puga et al., 2014; Wang et al., 2014). Notably, earlier studies indicated that inositol polyphosphates play a role in phosphate signaling, since defects in the INOSITOL PENTAKISPHOSPHATE 2-KINASE (IPK1), which catalyzes the conversion of InsP_5_ [2-OH] to InsP_6_, results in defective phosphate starvation responses (PSR) in Arabidopsis (Stevenson-Paulik et al., 2005; Kuo et al., 2014). In support of this, a more recent study showed that the inositol 1,3,4-trisphosphate 5-/6-kinase ITPK1 regulates phosphate homeostasis as well and that both *itpk1* and *ipk1* seedlings showed similar PSR phenotypes (Kuo et al., 2018). These phenotypes were proposed to be associated with increased levels of an InsP_4_ isomer of yet unknown isomer identity (Kuo et al., 2018). Work by Wild and colleagues (2016) pointed to a role of PP-InsPs in phosphate regulation, by showing that 5-InsP_7_ induces a physical interaction between the rice proteins OsPHR2 and OsSPX4 at micromolar concentrations. More recently, Arabidopsis mutants defective in the PPIP5K/Vip1-type InsP_7_ kinases VIH1 and VIH2, and hence defective in InsP_8_ synthesis, were shown to display disturbed PSR and strong phosphate over-accumulation, supporting the idea that PP-InsPs regulate phosphate homeostasis (Dong et al., 2019; Zhu et al., 2019).

Metabolic pathways leading to the production of PP-InsPs are well established in yeast and metazoan. There, IP6K/Kcs1-type kinases phosphorylate InsP_6_ at the 5 position, generating 5-InsP_7_ (Saiardi et al., 1999; Draskovic et al., 2008), while PPIP5K/Vip1 proteins catalyze the phosphorylation of 5-InsP_7_ to generate 1,5-InsP_8_ (Mulugu et al., 2007; Lin et al., 2009; Zhu et al., 2019). Vip1 enzymes are ubiquitously found in plants from green algae to monocot and eudicot angiosperms (Laha et al., 2015). However, the identity of enzymes responsible for InsP_7_ synthesis and roles for InsP_7_ in plants still remain elusive.

Recently, we demonstrated that recombinant Arabidopsis ITPK1 and ITPK2 phosphorylate InsP_6_ to 5-InsP_7_ *in vitro*, the major InsP_7_ isomer identified in Arabidopsis seeds (Laha et al., 2019). In this study, we characterized ITPK1-deficient plants in more detail and find that ITPK1-deficiency compromised inositol polyphosphate homeostasis, including reduced levels of InsP_7_ and InsP_8_. Our study reveals that *itpk1* plants display auxin perception phenotypes that are independent of disturbed PSR and demonstrate that a specific increase in 5-InsP_7_ potentiates the interaction of various TIR1 homologs with Aux/IAA-type transcriptional repressors in yeast. We also provide evidence for a direct interaction of ITPK1 with the auxin co-receptor component TIR1 *in planta*, suggesting a dedicated substrate channeling of ITPK1-dependent InsPs/PP-InsPs to activate auxin signaling.

## RESULTS

### ITPK1-deficient Plants Show Altered Inositol Polyphosphate Profile

The discovery that ITPK1 and ITPK2 exhibit InsP_6_-kinase activity *in vitro* (Laha et al., 2019) encouraged us to investigate the physiological function of these kinases. QPCR analyses revealed ubiquitous expression of *ITPK1* and *ITPK2* (Fig. 1A). While high levels of *ITPK1* transcripts were detected in all plant tissues, the expression of *ITPK2* was notably strong in the root and hypocotyl. To assess the contribution of ITPK1 and ITPK2 in InsP_7_ synthesis, we analyzed loss-of-function T-DNA insertion lines for both genes (Supplemental Fig. S1A). SAX-HPLC analyses of [^3^H]-inositol-labeled seedlings revealed no differences in InsP_7_ content between wild-type and the *itpk2-2* mutant (Supplemental Fig. S1B). Likewise, no differences in the amount of InsP_6_, the most abundant inositol phosphate, were found between Col-0 wild-type and *itpk1* seedlings (Fig. 1B-E and Supplemental Fig. S1C). However, *itpk1* seedlings displayed reduced levels of InsP_7_ and InsP_8_ (Fig. 1C, E), suggesting that ITPK1 functions as a cellular InsP_6_ kinase. Unchanged ratios of InsP_8_/InsP_7_ (Supplemental Fig. S1D) indicate that ITPK1 does not have InsP_7_ kinase function. In agreement/line with previously published *in vitro* data demonstrating that ITPK homologs can phosphorylate different InsP_3_ and InsP_4_ isomers (Sweetman et al., 2007; Stiles et al., 2008), we also observed compromised InsP_5_ [1/3-OH] levels and a strong increase in as yet unknown isomers of InsP_3_ and InsP_4_ in *itpk1* plants (Fig. 1B-C, E and Supplemental Fig. S1C). These observations are largely in agreement with recent findings of Kuo and colleagues (Kuo et al., 2018), who analyzed [^32^P]-labeled seedlings, except for a reduction in InsP_6_ in *itpk1* seedlings reported by these authors, which we did not detect with [^3^H]-inositol-labeling. To address this point in more detail, we analyzed unlabeled plants by TiO_2_ pulldown-based protocol with subsequent separation on PAGE and toluidine staining (Wilson et al., 2015). In agreement with our SAX-HPLC analyses of [^3^H]-inositol-labeled seedlings, we did not observe any differences in cellular InsP_6_ in *itpk1* seedlings (Fig. 1D, E).

**Figure 1.**
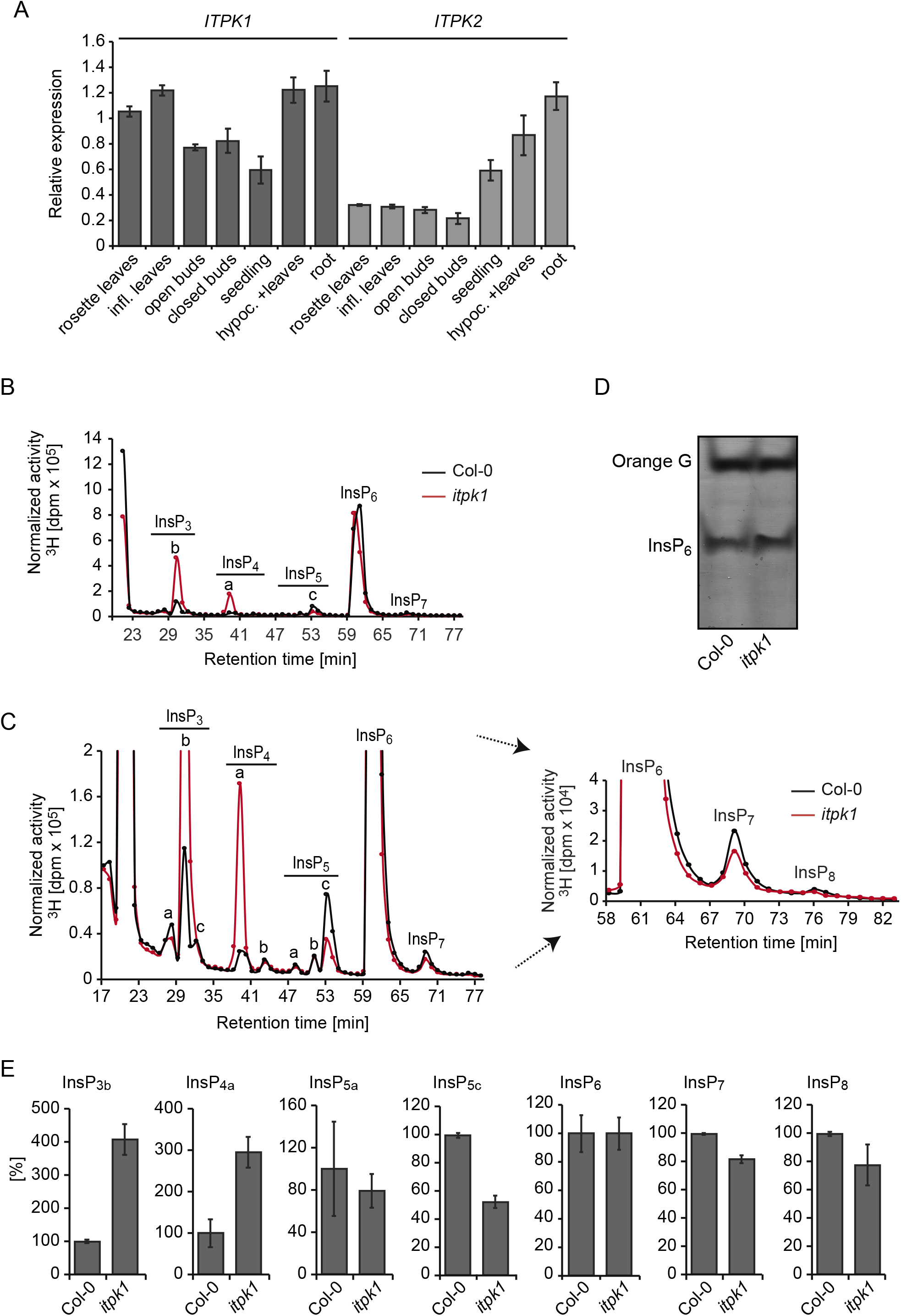
*ITPK1* and *ITPK2* are ubiquitously expressed and *itpk1* mutant plants display pleiotropic defects in inositol polyphosphate and inositol pyrophosphate homeostasis. **(A)** Expression analyses of Arabidopsis *ITPK1* and *ITPK2*. qPCR analyses of *ITPK1* and *ITPK2* using cDNA prepared from RNA extracts of different tissues of wild-type Col-0 plants. Error bars represent standard deviation (s.d.), n=3. *PP2AA3* was used as reference gene. **(B)** SAX-HPLC profiles of extracts of 3-week-old [^3^H] inositol-labeled wild-type (Col-0, solid black line) and *itpk1* mutant (solid red line) seedlings. Activities obtained by scintillation counting of fractions containing the InsP_2_-InsP_8_ peaks are shown. **(C)** A zoom in into the SAX-HPLC profiles of (B). The InsP_6_-InsP_8_ region is presented with arrows. The isomeric nature of InsP_3[a-c]_, InsP_4[a,b]_ is not yet solved. Based on published chromatographs (Stevenson-Paulik et al., 2005; Laha et al., 2015), InsP_5a_ corresponds to InsP_5_ [2-OH], InsP_5b_ represents InsP_5_ [4/6-OH] and InsP_5c_ corresponds to InsP_5_ [1-OH] or its enantiomer InsP_5_ [3-OH]. **(D)** Inositol polyphosphate enrichment from indicated plant extracts by TiO_2_ pulldown. Inositol polyphosphates were eluted from TiO_2_ beads, separated by PAGE and visualized by Toluidine blue. **(E)** Relative amounts of different inositol polyphosphates of 3-week old [^3^H] inositol-labeled seedlings. InsP_3b_, InsP_4a_, InsP_5a_, InsP_5c_ and InsP_8_ are presented as percentage of total inositol phosphates. InsP_6_ and InsP_7_ are presented as ratio to their precursor InsP_5a_. Error bars represent standard errors (s.e.m.), n=2. The experiment was repeated independently with similar results.

### ITPK1 Plays a Critical Role in Auxin-Related Processes

Next, we assessed the physiological consequences of the detected changes in inositol polyphosphates in *itpk1* plants. Compared to Col-0 wild-type, primary root elongation was impaired in *itpk1* plants (Supplemental Fig. S2A), a phenotype that could be fully rescued by introducing the genomic *ITPK1* fragment C-terminally fused to G3GFP (Supplemental Fig. S2A, S2B and S2C). This ITPK1-G3GFP fusion also rescued yeast *kcs1*Δ-associated growth defects (Supplemental Fig. S2D). We further noticed an *itpk1*-associated defect in leaf venation. The T-DNA insertional mutant displayed an increased number of end points, as well as compromised gravitropic root curvature (Fig. 2A and 2B; Supplemental Fig. S3A). Both defects were also robustly complemented in *itpk1* lines expressing the genomic *ITPK1* fragment.

**Figure 2.**
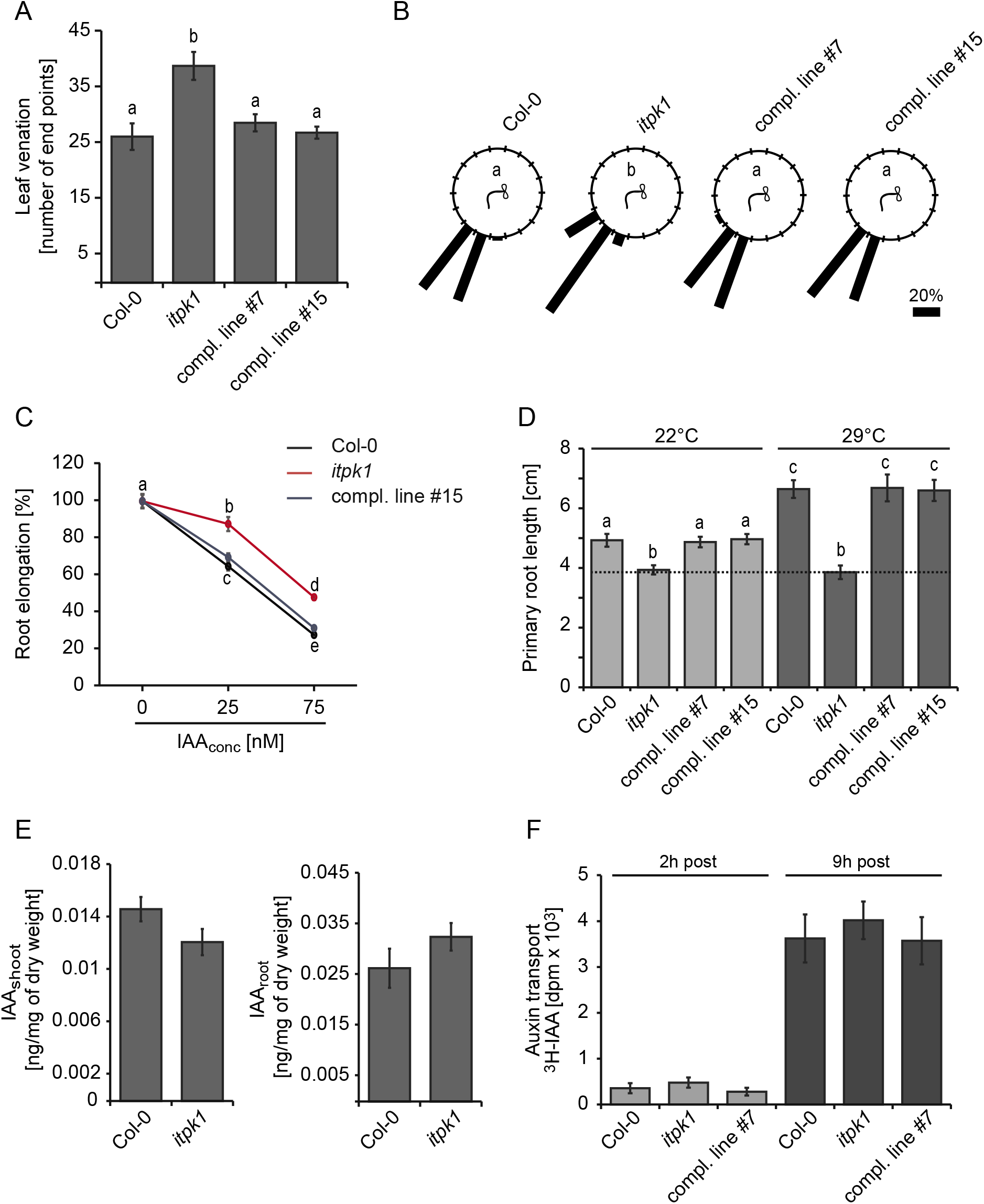
Loss of ITPK1 results in auxin signaling defects. **(A)** Analysis of leaf vascular differentiation. Venation patterns were recorded from 13-day-old seedlings of the indicated genotypes. Error bars depict s.e.m, n=6-10. The experiment was repeated twice with similar results. Letters depict significance in one-way analysis of variance (ANOVA) (a and b, *P* < 0.005). **(B)** Root gravitropism of 12-day-old seedlings. Indicated genotypes were rotated by 90° and gravitropic curvatures were measured after 16 h. The percentage of seedlings in each category are represented by the length of the bar. The distribution of data was analyzed using χ^2^ test (number of seedlings n ≥ 36, groups contained at least 7.5% of total seedlings per genotype). Significant differences (*P* <0.0001) are indicated by different letters. The experiment was performed independently with similar results. **(C)** Relative root elongation of designated genotypes under increasing IAA concentrations. 6-day-old seedlings were transferred to MS plates supplemented with 0, 25 and 75 nM IAA and incubated for 7 days. Root lengths were evaluated by ImageJ. Error bars are s.e.m, n=10-35. Different letters indicate significance in one-way analysis of variance (ANOVA) (a and b, *P* < 0.05; a to c, *P* < 0.001; a to d, *P* < 0.001; b to c, *P* < 0.001; d and e, *P* < 0.001; b to d, *P* < 0.001; c to e, *P* < 0.001). The experiment was repeated twice with similar results. **(D)** Primary root length analysis of designated genotypes grown at higher temperature. 5-day-old seedlings were kept at 22°C or shifted to 29°C. Root length was evaluated after 8 days by ImageJ. Error bars represent s.e.m, n ≥ 17. Letters depict significance in one-way ANOVA (a and b, *P* < 0.005; a to c, *P* < 0.001; b to c, *P* < 0.001). The experiment was repeated independently with similar results. **(E)** Auxin levels in shoot and roots of 2-week-old seedlings of designated genotypes grown on sterile MS media. Error bars represent s.e.m, n ≥ 7. The experiment was repeated with similar results. **(F)** Polar auxin transport. The apical end of excised stems of designated genotypes were placed in liquid MS media supplemented with [^3^H] IAA. After indicated times of incubation, the basal ends of the labelled stems were excised and the activity was determined by scintillation counting. Error bars represent s.e.m, n=3. Genotypes in all panels are as indicated. The term “compl. line” refers to the *itpk1* T-DNA insertion line transformed with a genomic fragment containing a 1839-bp region upstream of the *ITPK1* start codon in translational fusion with a C-terminal G3GFP.

The phenotypic defects exhibited by *itpk1* plants are reminiscent of impaired auxin signaling (Sieburth, 1999; Scarpella et al., 2006; Teale et al., 2006; Salehin et al., 2015; Weijers and Wagner, 2016). To further test this possibility, we performed auxin sensitivity assays (Lincoln et al., 1990; Ruegger et al., 1998) by determining the root length after exogenous application of the natural auxin indole-3-acetic acid (IAA). While wild-type seedlings displayed gradually shorter roots in the presence of increasing amounts of IAA, *itpk1* seedlings were more resistant to the exogenous supply of auxin (Fig. 2C; Supplemental Fig. S3B and S3C). Auxin-insensitivity of *itpk1* plants was fully rescued in independent complemented lines expressing the genomic *ITPK1* fragment (Fig. 2C; Supplemental Fig. S3C). We then examined thermomorphogenesis, an adaptation response to elevated temperatures controlled by auxin (Ruegger et al., 1998; Quint et al., 2016; Wang et al., 2016; Bellstaedt et al., 2019). As reported earlier, exposure of wild-type seedlings to 29°C resulted in increased primary root length (Wang et al., 2016). However, this high temperature-induced response was severely compromised in *itpk1* plants (Fig. 2D), resembling auxin receptor mutants (Wang et al., 2016). The reduced root development of *itpk1* plants under high temperature was rescued in independent complemented lines (Fig. 2D). With these observations, we concluded that *ITPK1* has an important function in auxin-related growth and developmental processes.

### Impaired Auxin Responses in *itpk1* Plants is Not Caused by Altered Auxin Synthesis or Transport

Auxin levels in shoots and roots, as well as polar auxin transport were similar between *itpk1* and Col-0 wild-type plants (Fig. 2E and 2F), suggesting that the observed phenotypic differences were not related to either the synthesis or the transport of auxin. We also tested sensitivity to the synthetic auxin analog 1-naphthaleneacetic acid (NAA), which can bypass auxin carrier-mediated transport mechanisms as it diffuses more easily through membranes than IAA (Delbarre et al., 1996). The *itpk1* line was also less sensitive to NAA when compared to wild-type plants (Supplemental Fig. S3D), confirming that altered auxin transport is unlikely to cause the observed phenotypes. In contrast, the expression of several marker genes associated with auxin signaling were compromised in *itpk1* plants and rescued in complemented lines (Supplemental Fig. S3E). Taken together, these results suggest that *itpk1* plants are defective in auxin responses.

### Defects in Phosphate Homeostasis and in Auxin Responses Are Largely Independent Consequences of *itpk1* Loss of Function

Recently, it was reported that *itpk1* plants exhibit constitutive PSRs, which result in phosphate over-accumulation in leaves when plants are grown with sufficient phosphate supply (Kuo et al., 2018). To assess whether increased phosphate accumulation can affect auxin sensitivity, we analyzed the *pho2* mutant, in which disrupted phosphate signaling downstream of PHR1 leads to phosphate over-accumulation (Aung et al., 2006; Bari et al., 2006). Notably, we did not find significant differences between Col-0 wild-type and *pho2-1* plants with respect to gravitropic responses, sensitivity of primary root growth to auxin or high temperature-induced primary root elongation (Supplemental Fig. S4). We also found that phosphorus over-accumulation in *itpk1* plants was largely unaffected by auxin (Supplemental Fig. S5A), suggesting that exogenously applied auxin does not alter phosphate accumulation in shoots. However, when plants were grown on low phosphate, a condition that strongly inhibits primary root elongation (Gutierrez-Alanis et al., 2018), the auxin insensitive primary root growth of *itpk1* plants was not observed anymore (Supplemental Fig. S5B and S5C). Overall, these findings indicate that the impaired PSR of *itpk1* plants under sufficient phosphate supply does not explain the auxin-related phenotypes and that both the uncontrolled phosphate accumulation and defective auxin responsiveness are independent responses.

### Defects in Auxin Responses Correlate with Altered Inositol Polyphosphate and Inositol Pyrophosphate Homeostasis

To dissect which inositol derivatives are possibly involved in auxin signaling, we employed different mutants affected in inositol polyphosphate synthesis. In line with the largely unchanged InsP profile, *itpk2-2* plants did not display defects in gravitropic responses nor did they show increased auxin sensitivity of primary root growth (Supplemental Fig. S6A and S6B). Similar to *itpk1*, auxin responses were also significantly decreased in *ipk1-1* mutant plants (Supplemental Fig. S6C and S6D), which have compromised conversion of InsP_5_ [2-OH] to InsP_6_ (Stevenson-Paulik et al., 2005; Kuo et al., 2014) and hence have reduced InsP_6_, InsP_7_ and InsP_8_ levels (Laha et al., 2015). Thus, common denominators between *ipk1-1* and *itpk1* mutant plants are decreases in InsP_5_ [1/3-OH], InsP_7_ and InsP_8_ and an increase in InsP_4a_ (Fig. 1B-E; Supplemental Fig. S1C and (Stevenson-Paulik et al., 2005; Laha et al., 2015), suggesting that one or several of these inositol polyphosphate isomers are critical for auxin signaling.

Next, we analyzed inositol polyphosphates in *itpk4-1* mutant plants. In agreement with recent findings of Kuo and colleagues (Kuo et al., 2018), we detected a robust reduction of InsP_6_ in *itpk4-1* as compared to wild-type Col-0 (Fig. 3A, 3B and 3C). We also observed reduced InsP_5_ [1/3-OH] levels in this mutant (Fig. 3B and 3C). However, in contrast to the findings by Kuo and colleagues (2018), our SAX-HPLC analysis of [^3^H]-inositol-labeled seedlings did not detect any changes in InsP_7_ and InsP_8_ levels in *itpk4-1* as compared to wild-type plants (Fig. 3B and 3C). Notably, root growth of the *itpk4-1* line was not compromised in auxin sensitivity (Fig. 3D). In conclusion, a global reduction in InsP_5_ [1/3-OH] or InsP_6_ does not appear to result in auxin-related phenotypes and *itpk1*-dependent defects in auxin perception seem to occur without decreasing InsP_3b_. In summary, these findings suggest that either the reduction in InsP_7_ or InsP_8_ or an increase in InsP_4a_, or a combinatorial derangement of these three inositol polyphosphates resulted in defective auxin responses in the *itpk1* and *ipk1-1* lines.

**Figure 3.**
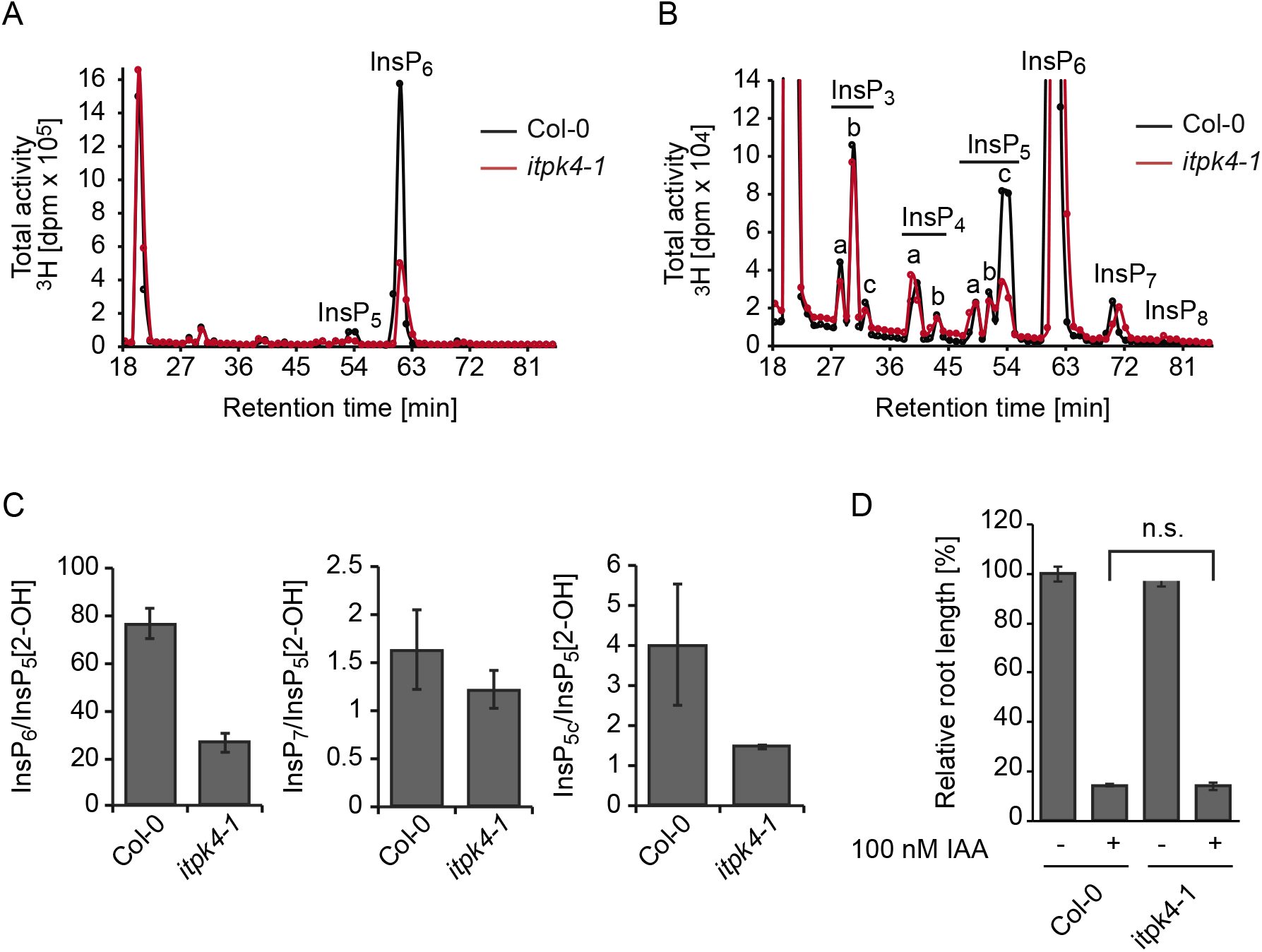
The *itpk4-1* plants are compromised in InsP_6_ but not in InsP_7_ synthesis and behave like wild-type Col-0 plants with respect to auxin-related processes. **(A)** SAX-HPLC profiles of extracts of [^3^H] inositol-labeled wild-type (Col-0, solid black line) and *itpk4-1* seedlings (solid red line). **(B)** Zoom-in into the SAX-HPLC profile of (A). **(C)** Relative amounts of different inositol polyphosphates in the respective genotypes are presented. Error bars represent ± s.e., n= 2. The experiment was repeated independently with similar results. **(D)** Relative root elongation of seedlings of designated genotypes treated with IAA. Seeds were surface sterilized and sown on sterile solid 0.5 x MS, 1 % sucrose media. After 2 days of stratification, germinated seedlings were allowed to grow for 6 days and transferred to solid MS media supplemented with or without 100 nM IAA. Root lengths were evaluated by ImageJ after another 5 days of growth. Data are means ± s.e., n=32-64.

To address the question of whether InsP_8_ is involved in auxin perception, we investigated two independent *vih2* lines (*vih2-3* and *vih2-4*), in which virtually no InsP_8_ can be detected in vegetative tissues (Laha et al., 2015). Notably, both mutants were undistinguishable from Col-0 wild-type plants with respect to thermomorphogenesis, gravitropism and sensitivity of the primary root to auxin (Supplemental Fig. S7A, S7B and S7C). Thus, these results suggest that bulk InsP_8_ has no critical role in auxin signaling.

### *In Vitro* Reconstitution Assays Suggest that Higher Inositol Polyphosphates Bind to the Auxin-receptor Complex with High Affinities

Considering the unknown isomer identity of the InsP_4_ species accumulating in *itpk1* lines (15 distinct InsP_4_ isomers are possible but only a few are commercially available, which allows limited evaluation of these isomers), we purified the InsP_4a_ species from the *itpk1* SAX-HPLC run. When incubated with recombinant ITPK1 and ATP, purified InsP_4a_ was efficiently phosphorylated (Supplemental Fig. S8A), suggesting that InsP_4a_ is an *in vivo* substrate of ITPK1. Furthermore, we performed direct auxin receptor complex binding assays with purified [^3^H]-InsP_3b_ and [^3^H]-InsP_4a_ that accumulate in ITPK1-deficient plants and found that binding of the [^3^H]-InsP_4_ species is detectible, although significantly reduced as compared to InsP_6_ (Supplemental Fig. S8B). No detectible binding of the [^3^H]-InsP_3_ species accumulating in *itpk1* seedlings was observed (Supplemental Fig. S8B). These data suggest that InsP_4a_ cannot be excluded as a potential direct (negative) regulator of auxin perception while InsP_3b_ is unlikely to play a direct role in auxin receptor regulation.

To further investigate the contribution of ITPK1 in auxin perception, we performed competitive binding assays with insect cell-purified ASK1-TIR1, recombinant Aux/IAA protein, natural auxin (IAA), and [^3^H]-InsP_6_ to determine IC_50_ values (50% displacement of radioligand binding) for ITPK1-dependent inositol polyphosphates and related molecules. IC_50_ values of different InsP species to compete with [^3^H]-InsP_6_ were as follows: InsP_6_ (IC_50_: 19 nM) ≤ 5-InsP_7_ (IC_50_: 20 nM) < InsP_5_ [3-OH] (IC_50_: 31 nM) < InsP_5_ [5-OH] (IC_50_: 34 nM) < InsP_5_ [1-OH] (IC_50_: 114 nM) (Fig. 4). These results indicated that the ASK1-TIR1-IAA complex has distinct binding affinities towards different inositol phosphate isomers (including enantiomers) with InsP_6_ and InsP_7_ displaying the highest affinities.

**Figure 4.**
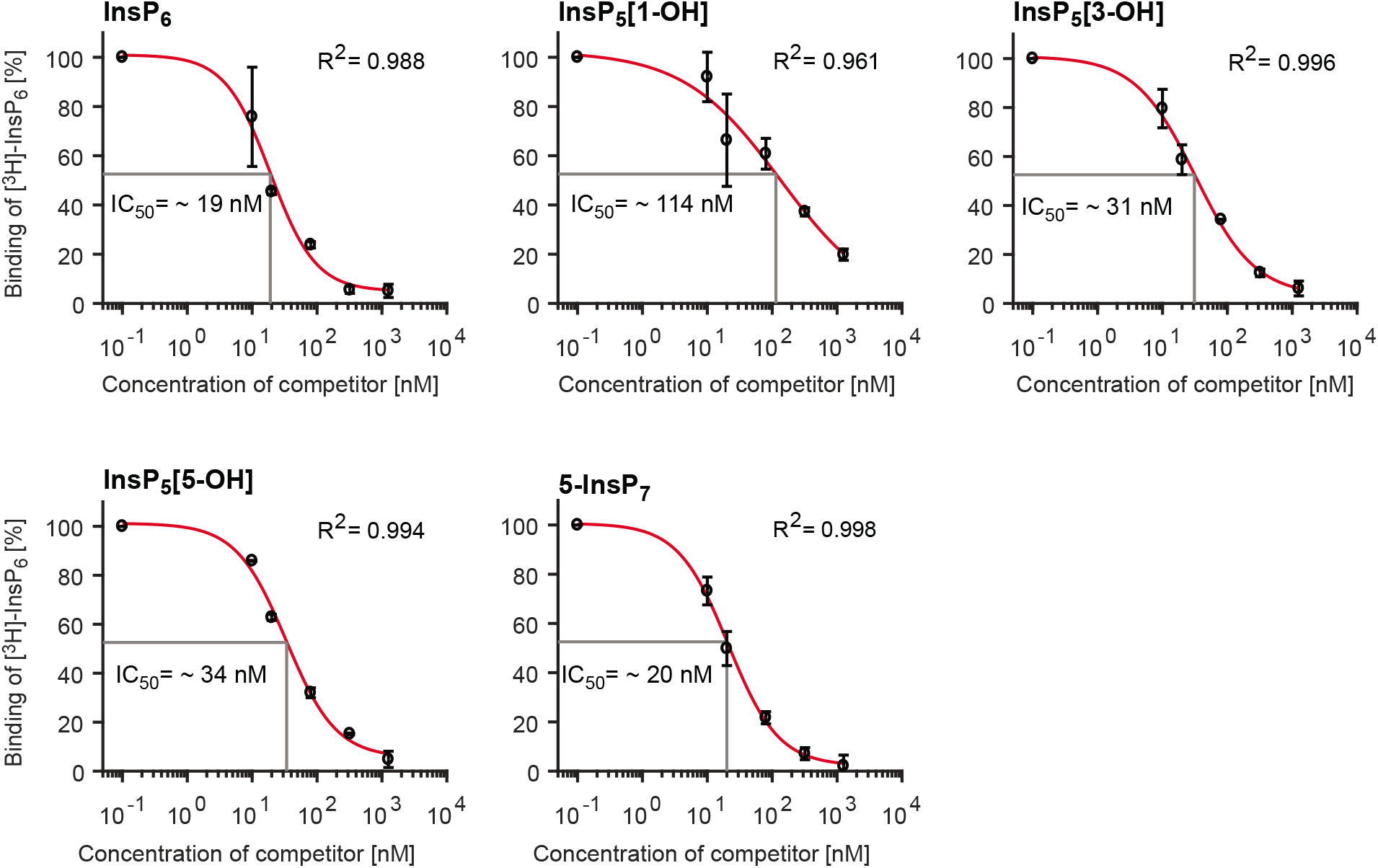
Higher inositol polyphosphates bind to the auxin receptor complex with higher affinities. Concentration-dependent competitive binding of [^3^H]-InsP_6_ to the ASK1-TIR1-Aux/IAA-IAA receptor complex in the presence of different unlabeled inositol polyphosphate species. Error bars represent s.e., n=2.

### The Inositol Pyrophosphate 5-InsP_7_ Potentiates the Interaction Between F-Box Proteins and Aux/IAA Repressors in Yeast

To delineate a potential role of 5-InsP_7_ in auxin-receptor complex formation *in vivo*, we performed yeast two-hybrid (Y2H) assays using the yeast EGY48 strain (Calderon Villalobos et al., 2012) and an isogenic strain lacking the *VIP1* gene. Mutation of *VIP1* in this yeast strain results in a specific accumulation of the 5-InsP_7_ isomer without changing levels of InsP_6_, InsP_4_ species, or any other inositol polyphosphate (Fig. 5A; Supplemental Fig. S9A), as also shown in other yeast genetic backgrounds (Onnebo and Saiardi, 2009; Laha et al., 2015). The interaction of the F-box protein AFB1 with the Aux/IAA proteins IAA5, IAA7, and IAA8 was robustly elevated in the *vip1*Δ strain (Fig. 5B). Similar results were also obtained when AFB2 was used as bait in the assay (Fig. 5C; Supplemental Fig. S9B), suggesting that 5-InsP_7_ potentiates auxin receptor complex formation in this system.

**Figure 5.**
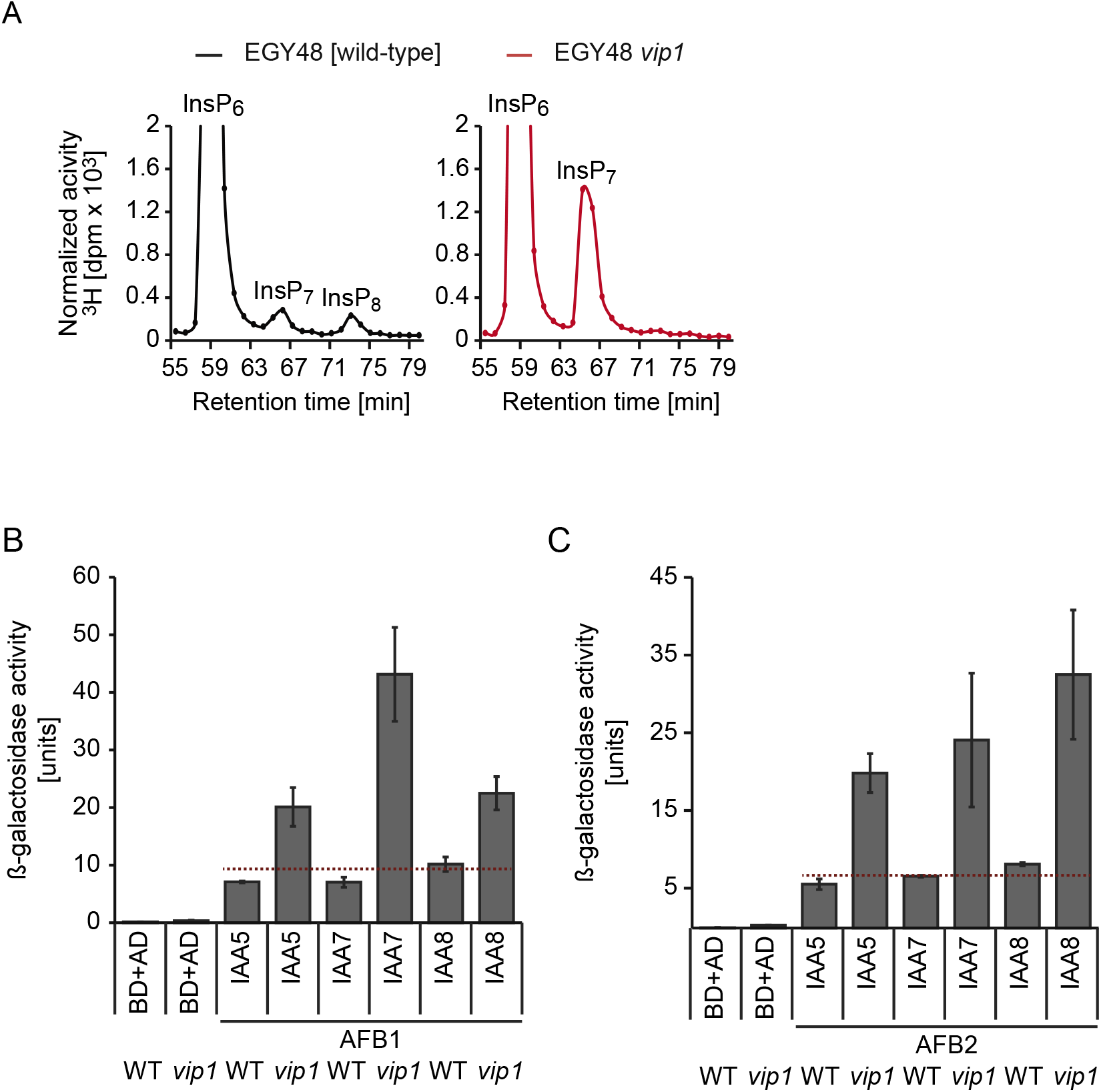
Inositol pyrophosphate 5-InsP_7_ activates auxin-receptor complex formation in yeast. **(A)** HPLC profiles of extracts of [^3^H] inositol-labeled wild-type and a *vip1*Δ yeast strains. Note that 5-InsP_7_ levels are specifically elevated in the *vip1*Δ yeast mutant. Extracts of designated strains were resolved by Partisphere SAX-HPLC. The full HPLC profile is presented in Supplemental Figure 9A. **(B, C)** Yeast two-hybrid (Y2H) assays to evaluate AFB1 and AFB2 interaction with different Aux/IAA repressors in isogenic *VIP1* and *vip*1Δ yeast strains. AFB1-Aux/IAA interactions in the presence of 1 μM IAA were quantified by β-galactosidase-mediated hydrolysis of ortho-nitrophenyl-b-D-galactopyranoside. Error bars depict s.e.m., n=3.

### ITPK1 Interacts with TIR1

A previously identified interaction of mammalian InsP_6_-kinase IP6K2 with a protein complex activated by casein kinase 2, an enzyme that requires InsP_7_ for full activity (Rao et al., 2014), suggested that PP-InsPs might be generated in close proximity to dedicated effector proteins. Since 5-InsP_7_, an ITPK1-dependent inositol pyrophosphate, binds with strong affinity to the TIR1-ASK1 receptor complex (Fig. 4), we investigated whether ITPK1 physically interacts with TIR1 to facilitate targeted delivery of 5-InsP_7_. To test this hypothesis, we first performed co-immunoprecipitation assays. We took advantage of Arabidopsis *itpk1* complemented with *gITPK1-G3GFP* complemented lines, for which expression of GFP-fused ITPK1 and endogenous TIR1 could be detected using GFP and TIR1 antibodies, respectively (Fig. 6A). Immunoprecipitation of ITPK1-G3GFP using GFP antibody allowed the subsequent detection of TIR1 as a co-immunoprecipitant, suggesting that ITPK1 associates with TIR1 *in vivo*.

**Figure 6.**
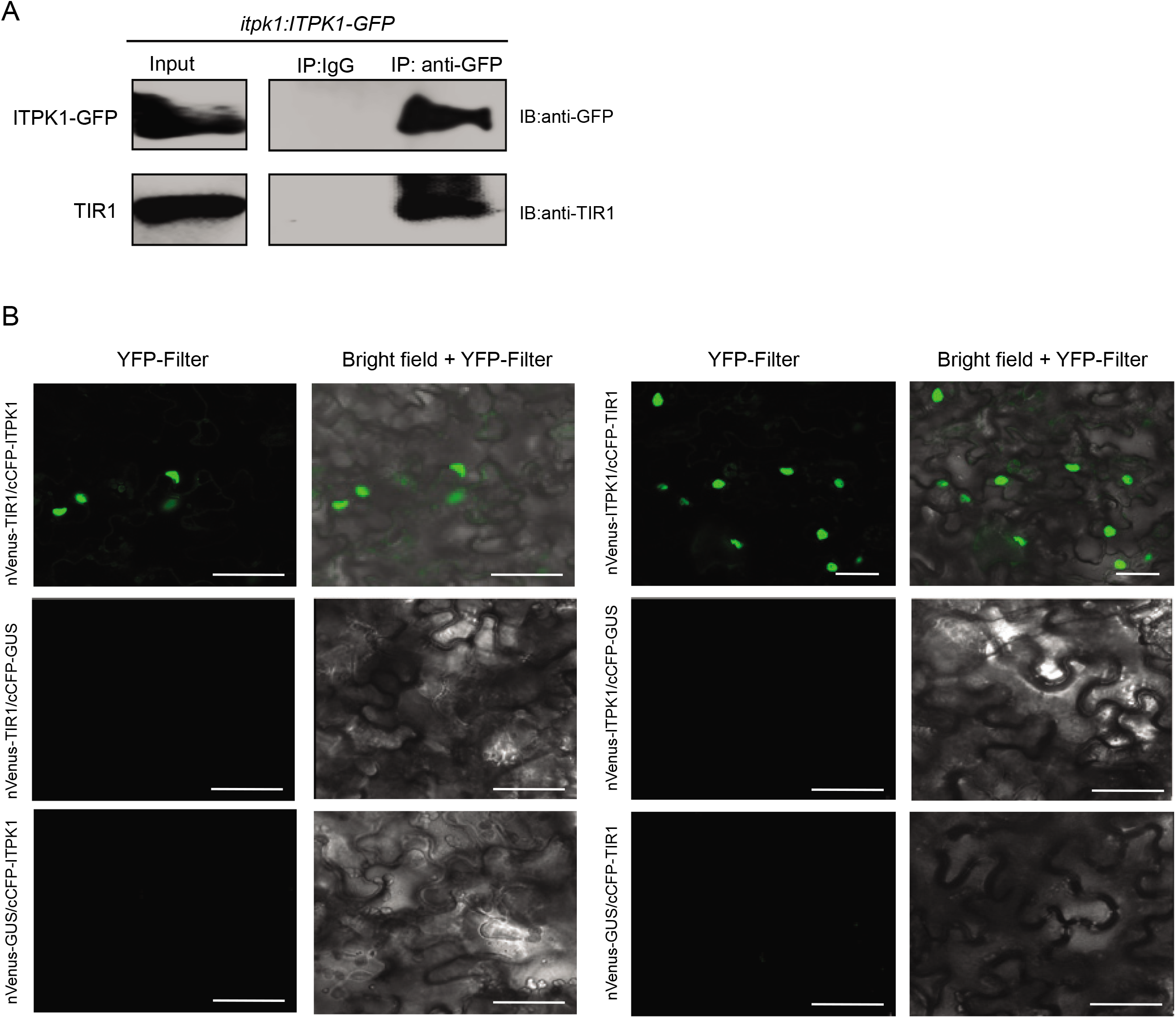
Arabidopsis ITPK1 physically interacts with TIR1. **(A)** Enrichment of TIR1 from *itpk1: ITPK1-G3GFP* transgenic plants. Total extracts from 4-week-old plants were immuno-enriched with either IgG- (control) or anti-GFP-conjugated agarose beads. Bound proteins were immunoblotted with anti-GFP or anti-TIR1 antibodies. Left panels indicate immunoblots of ITPK1-GFP or TIR1 in the input extracts. Right panel lanes are immunoblots of ITPK1-GFP and TIR1 for IgG- or anti-GFP immuno-enriched samples. **(B)** Transiently expressed ITPK1 interacts with TIR1 in the nucleus of *N. benthamiana* cells. Bi-molecular fluorescence complementation (BiFC) to detect interaction of ITPK1 with TIR1. Agrobacterium strains expressing indicated nVenus- or cCFP-tagged ITPK1, TIR1 or GUS (control) were combinatorially co-expressed in *N. benthamiana* leaves. At 2-days-post-infiltration (dpi), tissue sections were visualized under a laser confocal microscope. Interaction and nuclear localization of ITPK1 with TIR1 in reciprocal BiFC combinations is shown (top row). Lack of detectable YFP fluorescence in any BiFC combinations of ITPK1 or TIR1 with GUS (bottom two rows) is also shown. Images are presented as YFP fluorescence (YFP filter) and merge of bright field of the same section with YFP fluorescence (brightfield + YFP filter). The combinations of nVenus- and cCFP-expressing constructs for the corresponding images are shown in left. Scale bar = 50 μm.

To further validate the interaction between ITPK1 and TIR1, we used bimolecular fluorescence complementation (BiFC) assays. Both co-expressions of nVenus-ITPK1 with cCFP-TIR1 and of nVenus-TIR1 with cCFP-ITPK1 in *Nicotiana benthamiana* resulted in YFP fluorescence in the nucleus (Fig. 6B). In contrast, no fluorescence was detected in any BiFC combination of ITPK1 with GUS or of TIR1 with GUS, respectively (Fig. 6B). Taken together, these results demonstrate that ITPK1 physically interacts with TIR1.

## DISCUSSION

### Potential Mechanism of ITPK1-Dependent Auxin Perception

Recently, it has been shown that phosphate homeostasis is impaired in *ipk1-1* and *itpk1* plants (Kuo et al., 2014; Kuo et al., 2018). In the present study, we show that both mutants also exhibit various growth and developmental phenotypes associated with defective auxin perception. We also provide evidence that the auxin- and phosphate-related phenotypes of *itpk1* plants are largely independent, since: i) phosphate over-accumulation *per se* did not result in defective auxin response (Supplemental Fig. S4); and ii) additional auxin supply could not prevent uncontrolled phosphate accumulation in *itpk1* shoots (Supplemental Fig. S5). These results also reflect the distinct roles of different inositol polyphosphates, which are produced in an ITPK1-dependent manner (Fig. 1B-E).

The impaired auxin responses of *ipk1*-1 and *itpk1* plants are most likely not caused by compromised stability of the auxin co-receptor F-box protein TIR1, as suggested by immunoblot analyses (Supplemental Fig. S10A). Instead, we propose that ITPK1-dependent InsPs/PP-InsPs induce the auxin receptor complex, therefore resulting in more efficient auxin signaling. The previously published crystal structure of the auxin receptor complex with insect-purified InsP_6_ (Tan et al., 2007) revealed a large hydrophilic binding pocket in which InsP_6_ is coordinated by extensive interactions with basic residues of the concave surface of the TIR1 leucine rich repeat (LRR) solenoid (Supplemental Fig. S10B) (Tan et al., 2007). Reminiscent for what has been predicted for the InsP_8_-bound F-box component COI1 of the jasmonate receptor complex (Laha et al., 2016), these interactions are highly anisotropic. As illustrated in Supplemental Fig. S10B, basic residues distal and proximal to the hormone binding pocket likely contribute to the elliptical shape of the LRR solenoid. Notably, the 5-position of InsP_6_ is protruding towards the IAA7 degron residue R90. The distance between the 5-phosphate and closest amino group of the R90 side chain of 5.6 Å suggests that the pyrophosphate moiety of 5-InsP_7_, if oriented similarly as InsP_6_, would stabilize the IAA7 degron interaction with an additional strong polar interaction. Considering the architecture of the SCF^TIR1^ auxin receptor complex (Dinesh et al., 2016), even small changes in the orientation of the IAA degron relative to TIR1 are likely to have strong consequences with respect to the ubiquitination efficiency and thus degradation of the Aux/IAA repressor and subsequent activation of auxin-responsive gene expression. In absence of a 5-InsP_7_-bound crystal structure of the auxin receptor complex, *in silico* docking experiments combined with molecular dynamics simulations and in parallel *cryo* EM studies could provide more mechanistic details on the role of 5-InsP_7_ on auxin receptor function.

Our results point to a possible direct role of ITPK1-dependent inositol polyphosphates in auxin perception. This idea was corroborated by yeast two-hybrid assays that suggest 5-InsP_7_ might represent a critical co-ligand of the ASK1-TIR1-Aux/IAA auxin co-receptor complex controlling auxin perception. However, we cannot exclude the possibility that other species altered in *itpk1* plants (or combinations thereof), such as InsP_5_ [1/3-OH] or isomers of InsP_3_, InsP_4_ and InsP_8_ may also contribute to the auxin-perception defect in these plants. The concept that efficient auxin perception by the auxin receptor complex might require two ligands, i.e., auxin and 5-InsP_7_ or another ITPK1-dependent inositol polyphosphate, appears reminiscent of jasmonate perception reported to rely on the coincidence detection of a JA-conjugate and the inositol pyrophosphate InsP_8_ (Laha et al., 2015). Such coincidence detection might be used to fine-tune hormone response depending on other internal or external cues.

### Substrate Channeling by ITPK1

Our work reveals a physical interaction of ITPK1 with TIR1 (Fig. 6). While further work will be necessary to establish whether this interaction is of functional relevance and how it is regulated, we hypothesize that the apparent close proximity of ITPK1 to the auxin receptor complex might create a privileged, local enrichment of InsPs/PP-InsPs to a dedicated effector protein complex similar to what has been observed for casein kinase 2 in mammalian cells (Rao et al., 2014). While we cannot exclude the possibility that the interaction of ITPK1 and TIR1 might help to generate an important signaling molecule at minimal energetic cost, we speculate that this interaction more likely represents a mechanism to prevent or to control the stimulation of other signaling events triggered by the same molecule. This would be particularly intriguing for PP-InsPs, whose long distance cellular movement is likely restricted, assuming their lifetime is as similarly short as mammalian PP-InsPs (Menniti et al., 1993).

To better understand the role of 5-InsP_7_ and other ITPK1-dependent InsPs and/or PP-InsPs in auxin perception, it will be important to examine whether also AFBs interact with ITPK1 or other inositol polyphosphate kinases. It will also be important to know whether such dedicated interactions, and hence localized generation of regulatory molecules, might contribute to the specificity by which auxin controls many different aspects of plant growth and development. To explore if and how InsPs and PP-InsPs can contribute to the crosstalk between jasmonate and auxin signaling, and nutritional cues, tools have to be developed to follow InsPs and PP-InsPs with high specificity and with high temporal and spatial resolution. The work presented here provides a first step to unveil the physiological processes regulated by 5-InsP_7_ and other ITPK1-dependent InsPs and PP-InsPs and opens new avenues to manipulate and better understand inositol pyrophosphate signaling in eukaryotic cells.

## METHODS

### Plant Material and Growth Conditions

Seeds of T-DNA insertion lines of *Arabidopsis thaliana* (ecotype Col-0) were obtained from The European Arabidopsis Stock Centre (http://arabidopsis.info/). *The itpk1* (SAIL_65_D03), *itpk 2-2* (SAIL_1182_E03) and *itpk4-1* (SAIL_33_G08) were genotyped for homozygosity using T-DNA left and right border primers and gene-specific sense or antisense primers (Table S1). The *pITPK1:gITPK1-G3GFP* construct was generated as described in the cloning section. Auxin assays were performed with *vih2-3, vih2-4*, and *ipk1-1* (Laha et al., 2015), as well as *pho2-1* (Delhaize and Randall, 1995) mutant plants. Wild-type Col-0 and all relevant transgenic lines were amplified together on a peat-based substrate (GS90) under identical conditions (16 h light and 8 h dark, day/night temperatures 22/18°C and 120 μmol^−1^ m^−2^ light intensity), and seeds of the respective last progenies were harvested and used for all analyses described in this article. For sterile growth, seeds were surface sterilized in 70% (v/v) ethanol and 0.05% (v/v) Triton X-100 for 30 min and washed twice with 90% (v/v) ethanol. Sterilized seeds were sown on solid plant media supplemented with 0.5x MS (Murashige and Skoog), 1% sucrose, 0.8% phytagel, stratified for 2 days at 4°C, and grown under conditions of 8 h light (23°C) and 16 h dark (21°C), unless mentioned otherwise.

### Constructs and Strains

The *IAA7* ORF was amplified from cDNA of 2-week-old wild-type Col-0 seedlings. The forward and reverse primes used to amplify the ORFs contained restriction sites as indicated in Table S1. Amplified PCR products were inserted into CloneJET™ (Thermo Scientific) following the manufacturer’s instructions. The ORFs were then excised from respective CloneJET™ vectors and subcloned into the pET28-His_8_-MBP vector (Laha et al., 2015).

For complementation of *itpk1* plants, a genomic fragment encoding ITPK1 including a 1839-bp region upstream of the *ITPK1* start codon was amplified from wild-type Col-0 genomic DNA, cloned into pENTR-D-TOPO and recombined with pGWB550 (Nakagawa et al., 2007) to generate a plant transformation vector containing the genomic *ITPK1* in translational fusion with a C-terminal G3GFP. Transformed lines were selected on solid plant media with 25 μg/mL hygromycin.

For yeast two-hybrid assays, the EGY48 *vip1*Δ strain was generated following a strategy as previously described (Hegemann and Heick, 2011). Briefly, a marker gene cassette flanked by *loxP* sites was amplified from pUG6 (Euroscarf) with primers harboring 40 nucleotide-5’-overhangs for homologous recombination, as listed in Table S1. The DDY1810 strain was transformed with the PCR products. Knockout strains were genotyped with gene promoter-specific forward and cassette-specific reverse primers. Yeast transformation was performed by the Li-acetate method (Gietz et al., 1992).

### Extraction and SAX-HPLC Analyses of Yeast and *A. thaliana*

Extraction and quantification of inositol polyphosphates from yeast and from Arabidopsis seedlings were carried out as described (Laha et al., 2015). For the latter, 10-day-old Arabidopsis seedlings grown in 8 h light/16 h dark condition in media supplemented with 0.5x MS, 1% sucrose, 0.8% phytagel were transferred to 3 mL liquid sterile media containing 0.5x MS, pH 5.7, 30 μCi mL^−1^ of [^3^H]-*myo*-inositol (30 to 80 Ci mmol^−1^; Biotrend; ART-0261-5). Seedlings were allowed to label for 6 days, then washed twice in dH2O and frozen in liquid N2. Extraction of inositol polyphosphates was done as described (Azevedo and Saiardi, 2006) and extracts were resolved by strong anion exchange high performance liquid chromatography (SAX-HPLC) using a Partisphere SAX 4.6 x 125 mm column (Whatman) at a flow rate of 0.5 mL min^−1^ with a shallow gradient formed by buffers A (1 mM EDTA) and B [1 mM EDTA and 1.3 M (NH4)2HPO4, pH 3.8, with H3PO4] (Laha et al., 2015).

### Root Gravitropism Assays

Seedlings were grown on vertical plates containing sterile 0.5x MS media in 8 h light/16 h dark condition. After 7 days, seedlings were transferred to solid plant media supplemented with 0.5x MS, 1% sucrose and 0.8% phytagel. Unless mentioned otherwise, after another 7 days of growth, the seedlings of indicated genotypes were rotated by 90° and the gravitropic curvature was measured at different time points and scored in categories of 20°.

### Leaf Venation Analyses

Fixation and dehydration of plant tissue was done as described (Sieburth, 1999). Briefly, the plant tissue was immersed overnight in a 3:1 mixture of ethanol: acetic acid and then dehydrated through 80%, 90%, 95% and 100% (v/v) ethanol. For clearing, the tissue was incubated in saturated chloral hydrate solution (2.5 g/mL) overnight. The tissue was visualized and imaged by a Zeiss Axio zoomV16 microscope system. Leaf venation analyses were performed using LIMANI (Dhondt et al., 2012).

### Auxin Transport

Polar auxin transport assays were performed as previously described (Ruegger et al., 1998). In short, 4 cm long inflorescence stems of 6-week-old plants were excised and the apical end was placed into a 1.5 mL microfuge tube containing 50 μL of labelling solution supplemented with 0.5x MS, 1% sucrose, pH 5.7 and 0.06 nCi mL^−1^ of [^3^H]-indole-3-acetic acid (15 to 30 Ci mmol^−1^; Biotrend; ART 0340). The excised stems were incubated in the solution for different time points. A 0.8-cm-section from the basal end of the labelled stem was excised and placed in a vial containing 2 mL of scintillation cocktail and incubated overnight before radioactivity was measured by scintillation counting.

### Hormone Measurements

Roots and shoots of 2-week-old seedlings grown in 8 h light/16 h dark condition were excised and collected separately. Auxin measurements were done as previously described (Eggert and von Wiren, 2017).

### Gene Expression Analyses

RNA extraction, cDNA synthesis and qPCR analyses were performed as previously described (Laha et al., 2015). Seedlings were grown vertically on sterile media supplemented with 0.5x MS, 1% sucrose, 0.8% phytagel for 11 days in 8 h light/16 h dark condition, then transferred to liquid media supplemented with 0.5x MS, 1% sucrose, pH 5.7 and allowed to grow for another 2 days before harvesting. Total RNA extraction was performed using the RNeasy Plant Mini Kit (Qiagen). For cDNA synthesis, 1 μg of RNA was treated with DNase I. The reverse transcription was performed according to the manufacturer’s instructions (Roboklon; AMV Reverse Transcriptase Native). SYBR Green reaction mix (Bioline; Sensimix SYBR No-ROX kit) was used in a Bio-Rad CFX384 real-time system for qPCR. The results were evaluated using the Bio-Rad CFX Manager 2.0 (admin) system. *PP2AA3* was used as a reference gene.

### *In Vitro* Kinase Assay

Recombinant enzymes were purified as described for the yeast Sfh1 protein (Schaaf et al., 2006). InsP_6_ kinase assays were performed by incubating enzymes in 15 μL reaction volume containing 20 mM HEPES (pH 7.5), 5 mM MgCl2, 5 mM phosphocreatine, 0.33 units creatine kinase, 12.5 mM ATP, 1 mM InsP_6_ and 1 mM DTT. The reaction was incubated at 28°C for 4 h, separated by PAGE and stained by toluidine blue (Losito et al., 2009). The InsP_4a_ kinase assay with recombinant ITPK1 was performed using the same reaction conditions described above. Here, InsP_4a_ was purified from [^3^H]-inositol labelled *itpk1* plants as described earlier (Blüher et al., 2017).

### *In Vitro* Radioligand-Binding-Based Reconstitution Assays

The *in vitro* binding assays were performed as previously described (Laha et al., 2016). InsP_5_ isomers were obtained from Sichem (Bremen, Germany) and [^3^H]-InsP_6_ was purchased from Biotrend (ART-1915-10). His_8_-MBP-IAA7 was isolated using a protocol used for protein purification of the yeast Sfh1 (Schaaf et al., 2006). ASK1-TIR1 was purified from insect cells as described (Tan et al., 2007). Insect cell-purified ASK1-TIR1 was incubated with recombinant His_8_-MBP-IAA7 at a molar ratio of 1:3 and in presence of 0.1 μM indole-3-acetic acid. [^3^H]-InsP_6_ was added to the binding buffer consisting of 50 mM Tris-HCl, pH 7.5, 100 mM NaCl, 10 mM imidazole, 10% (v/v) glycerol, 0.1% (v/v) Tween 20, and 5 mM 2-mercaptoethanol (freshly added) in a total volume of 0.5 mL. The reaction mixture was then incubated at 4°C for 2 h. Then 30 μL of Ni-NTA resin was added, the reaction was vortexed briefly and incubated further at 4°C for 3 h. A total of 10 mL ice-cold washing buffer (reaction buffer without 2-mercaptoethanol) was added to individual reactions and after 20 sec Ni-NTA beads were filtered with 2.4-μm glass fiber filters (25 mm; Whatman Cat No 1822 025) using a filtration system (model FH225V, Hoefer, San Francisco), before analyzing filter membranes by scintillation counting. Data collection and evaluation of IC_50_ was carried out as described (Laha et al., 2016).

### Yeast Two-Hybrid Assays

The *Saccharomyces cerevisiae* yeast strain EGY48 (p8opLacZ) (Sheard et al., 2010) was transformed with pGILDA-*AFB1* or pGILDA-*AFB2* together with *pB42AD-IAA5*, pB42AD-*IAA5* or *pB42AD-IAA8* constructs (Calderon Villalobos et al., 2012), following the standard yeast transformation protocol (Gietz et al., 1992). Yeast transformants were selected on solid CSM media with appropriate supplements lacking uracil, tryptophan and histidine, and spotted in 8-fold serial dilutions onto CSM media with appropriate supplements lacking uracil, tryptophan, histidine and leucine. For quantification, fresh colonies from independent transformants were grown overnight in liquid CSM media with appropriate supplements lacking uracil, tryptophan and histidine. Aux/IAA interactions with AFB1 or 2 in the presence of 1 μM indole-3-acetic acid were evaluated by quantification of β-galactosidase-mediated hydrolysis of ortho-nitrophenyl-β-D-galactopyranoside.

### Protein Extraction and Immunoblots from *Arabidopsis thaliana*

For the detection of endogenous TIR1 levels, total proteins were extracted in 6M Urea from 2-week-old indicated Arabidopsis plants. The extract was centrifuged at 15,000 x g for 10 min at 4°C. The supernatant was mixed with Laemmli loading dye to final 1x. The proteins were separated on a 7.5% SDS-PAGE, electroblotted onto a PVDF membrane and immunoblot was performed with anti-TIR1 (Agrisera) or anti-actin (Abiocode) antibodies. Blots were developed by ECL kit (Biorad) and images acquired using the Alpha imagequant system (GE).

For co-immunoprecipitation assays, 3-week-old *itpk1:ITPK1-G3GFP* transgenic plants were used. Leaf tissues were homogenized (w/v) in RIPA buffer (5 mM Tris-HCl pH 7.5, 150 mM NaCl, 10mM MgCl2, 1 mM EDTA, 1% NP-40, 1 mM Sodium deoxycholate) containing 1X plant protease inhibitors (Sigma). The supernatant was clarified through a fine mesh (100 μm) and centrifuged at 5000 x g for 10 min at 4°C. The supernatant was then pre-cleared for 1 h with 25 μL of IgG agarose beads (Sigma) at 4°C with constant rotation. The beads were collected by centrifugation and washed three times with RIPA buffer. To the supernatant 25 μL of anti-GFP agarose beads (Biobharati) were added and kept at rotation overnight at 4°C. On the following day, the anti-GFP agarose beads were washed as described for the IgG-beads. Both IgG- and anti-GFP-agarose beads were resuspended in 100 μL of 1x Laemmli loading dye and separated on 7.5% SDS-PAGE gel. After transfer to a PVDF membrane and blocking, immunoblots were performed with anti-GFP (Biobharti) or anti-TIR1 antibodies (Agrisera). Images were acquired as described above.

### Constructs for Transient Expression, BiFC and Co-Immunoprecipitation of Proteins from *N. benthamiana*

*ITPK1* and *TIR1* sequences were amplified from total cDNA from wild-type Col-0 plants. The primer sequences are shown in Table 1. The amplicons were first cloned into the entry vector pDONR207 (Thermofisher) using BP Clonase (Invitrogen) and subsequently recombined using Clonase LR (Invitrogen) into the destination vectors pMDC43 (GFP) (Curtis and Grossniklaus et al., 2003), HA-pBA, pMDC-nVenus or pMDC-cCFP (Bhattacharjee et al., 2011), as indicated. The pENTR-GUS plasmid (Invitrogen) was similarly cloned into the BiFC destination vectors pMDC-nVenus or pMDC-cCFP. Resulting clones were confirmed by sequencing. The binary vectors were electroporated into the agrobacterium strain GV3101.

For BiFC assays, the indicated agrobacterium strains were cultured overnight in LB nutrient broth containing appropriate antibiotics. The cultures were then pelleted and resuspended in equal volumes of induction medium (10 mM MgCl_2_, 10 mM MES, pH 5.6) also containing 100 μM acetosyringone (Sigma) and maintained at room temperature for 4 h. Indicated combinations of agrobacterium suspensions were mixed at similar cell densities and infiltrated into *N. benthamiana* leaves. About 48 hours post infiltration, tissue sections from the infiltrated leaves were viewed under a SP8 confocal microscope (Leica-microsystems). Images were acquired with both bright field and YFP filters.

### Elemental Analysis

Whole shoot samples were dried for 48 h at 65°C and digested with nitric acid in polytetrafluoroethylene vials in a pressurized microwave digestion system (UltraCLAVE IV, MLS GmbH). Total phosphorus concentrations were analyzed by sector-field high-resolution inductively coupled plasma mass spectrometry (HR-ICP-MS; ELEMENT 2, Thermo Fisher Scientific, Germany). Element standards were prepared from certified reference standards from CPI-International (USA).

### Statistical Analysis

One-way ANOVA analysis followed by a Dunnett’s post hoc test on SPSS 24 (IBM), Chi-squared test and Student’s *t*-test were applied for statistical analysis.

### Accession Numbers

Sequence data from this article can be found in the GenBank/EMBL databases under the following accession numbers: *ITPK1* (At5g16760), *ITPK2* (At4g33770), *ITPK4* (At2G43980), *PP2AA3* (At1g13320), *IAA19* (At3g15540), *IAA5* (At1g15580), *ARF19* (At1g19220), *IAA29* (At4g32280), *TIR1* (At3g62980) and *β-TUBULIN* (At5g62700).

## Supplemental Data

The following supplemental materials are available.

**Supplemental Figure S1**. Inositol polyphosphate analyses of Arabidopsis *ITPK1* and *ITPK2* T-DNA insertion lines.

**Supplemental Figure S2**. ITPK1 regulates seedling growth and development in Arabidopsis and a C-terminal GFP Fusion of ITPK1 does not compromise ITPK1 functions.

**Supplemental Figure S3**. Plant responses to exogenous auxin are regulated by ITPK1.

**Supplemental Figure S4**. Auxin-related growth and developmental processes are not affected in Arabidopsis *pho2-1* Plants.

**Supplemental Figure S5**. Role of ITPK1 in phosphorus accumulation and the effect of auxin.

**Supplemental Figure S6**. The *ipk1-1* but not *itpk2-2* mutant is defective in auxin perception.

**Supplemental Figure S7**. VIH2-deficient plants are not compromised in auxin perception.

**Supplemental Figure S8**. Binding of inositol polyphosphates to the auxin-receptor complex.

**Supplemental Figure S9**. 5-InsP_7_ potentiates formation of auxin receptor complex *in vivo*.

**Supplemental Figure S10**. Stability of TIR1 in *itpk1* plants and structural considerations of inositol polyphosphate binding of the auxin receptor complex.

**Supplemental Table 1**. Primer list

**Supplemental References**.

## ACKNOWLEDGEMENTS

We thank Anna. L. Keller, Doris Pätzold, Sven T. Bitters, Brigitte Überbach, and Li Schlüter for technical assistance, Dr. Kai Eggert for phytohormone analysis and Marília Kamleitner for helpful suggestions and critically reading. We thank Tsuyoshi Nakagawa for Gateway binary vectors. We are grateful to Claudia Oecking for providing us with ^3^H-IAA. We are also thankful to Sheng Yang He, John Withers and Luz Irina Calderón Villalobos for providing us with the yeast EGY48 strain and several pGILDA, pB42AD constructs. This work was funded by grants from the Deutsche Forschungsgemeinschaft (SCHA 1274/4-1, SCHA 1274/5-1, Research Training Group GRK 2064 and under Germany’s Excellence Strategy – EXC 2070 – 390732324, PhenoRob to G.S.; and CIBSS – EXC 2189 – 390939984 as well as JE 572/4-1 to H.J.J.). D.L. acknowledges the Deutsche Forschungsgemeinschaft (LA 4541/1-1) for a Research Fellowship. H.G. and Y.W.D. are supported by DBT-JRF and DBT-YI fellowships, respectively. S.B. is supported by funds from DBT-Ramalingaswami Re-Entry Fellowship. A.S. is supported by the MRC/UCL Laboratory for Molecular Cell Biology University Unit (MC_UU_12018/4). Z.H.S. was supported by a scholarship from the Iran Ministry of Science, Research and Technology. N.Z. is a Howard Hughes Medical Institute Investigator.

The authors declare no competing interests.

