## Supplemental Data for "ITPK1-Dependent Inositol Polyphosphates Regulate Auxin Responses in *Arabidopsis thaliana*"

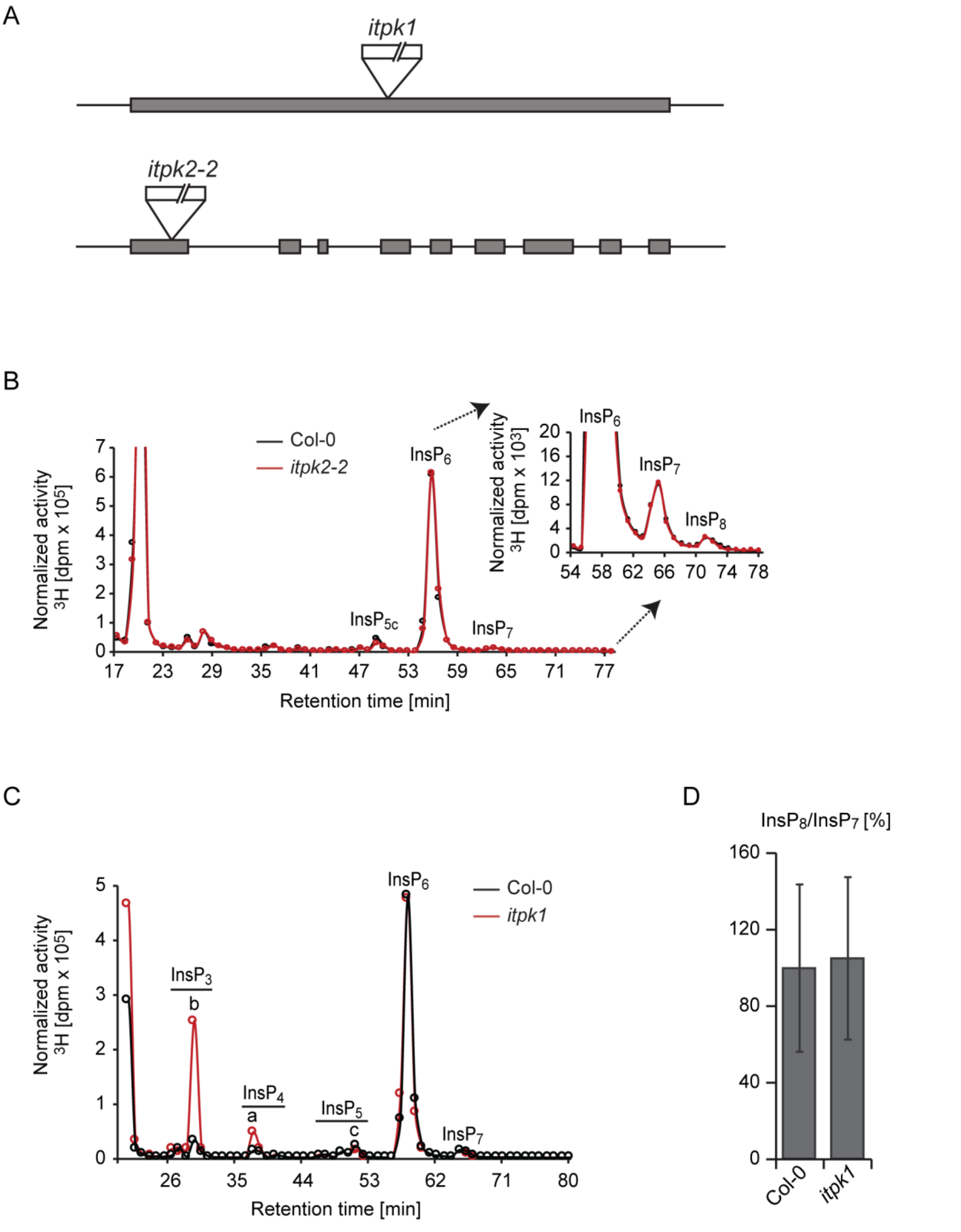


**Supplemental Fig. S1**. Inositol polyphosphate analyses of Arabidopsis *ITPK1* and *ITPK2* T-DNA insertion lines.

**(A)** Cartoon depicting the genome structure of *ITPK1* and *ITPK2* highlighting the position of T-DNA insertions in *itpk1* and *itpk2-2* lines.
**(B)** The *itpk2-2* line appears not to be compromised in inositol polyphosphate synthesis. Extracts of designated [$^3$H] inositol-labeled Arabidopsis seedlings were resolved by SAX-HPLC. Activities obtained by scintillation counting of fractions containing the $InsP_2$-$InsP_8$ peaks are presented.
**(C)** SAX-HPLC profiles of extracts of 3-week old [$^3$H] inositol-labeled Col-0 and *itpk1* seedlings. Activities obtained by scintillation counting of fractions containing the $InsP_2$-$InsP_8$ peaks are shown. HPLC runs from independent experiments are shown in Figure 5B. The isomeric nature of $InsP_{3[a\text{-}c]}$, $InsP_{4\,[a,b]}$ is not yet solved. Based on published chromatographs (Stevenson-Paulik et al., 2005; Laha et al., 2015), $InsP_{5a}$ corresponds to $InsP_5$ [2-OH], $InsP_{5b}$ represents $InsP_5$ [4/6-OH] and $InsP_{5c}$ corresponds to $InsP_5$ [1-OH] or its enantiomer $InsP_5$ [3-OH]. Experiments were repeated independently with similar results.
**(D)** Relative amounts of inositol polyphosphates of 3-week old [$^3$H] inositol-labeled seedlings. Data are presented as $InsP_8$/$InsP_7$ ratio (a measure of Vip1/PPIP5K activity). Error bars represent standard errors (s.e.).

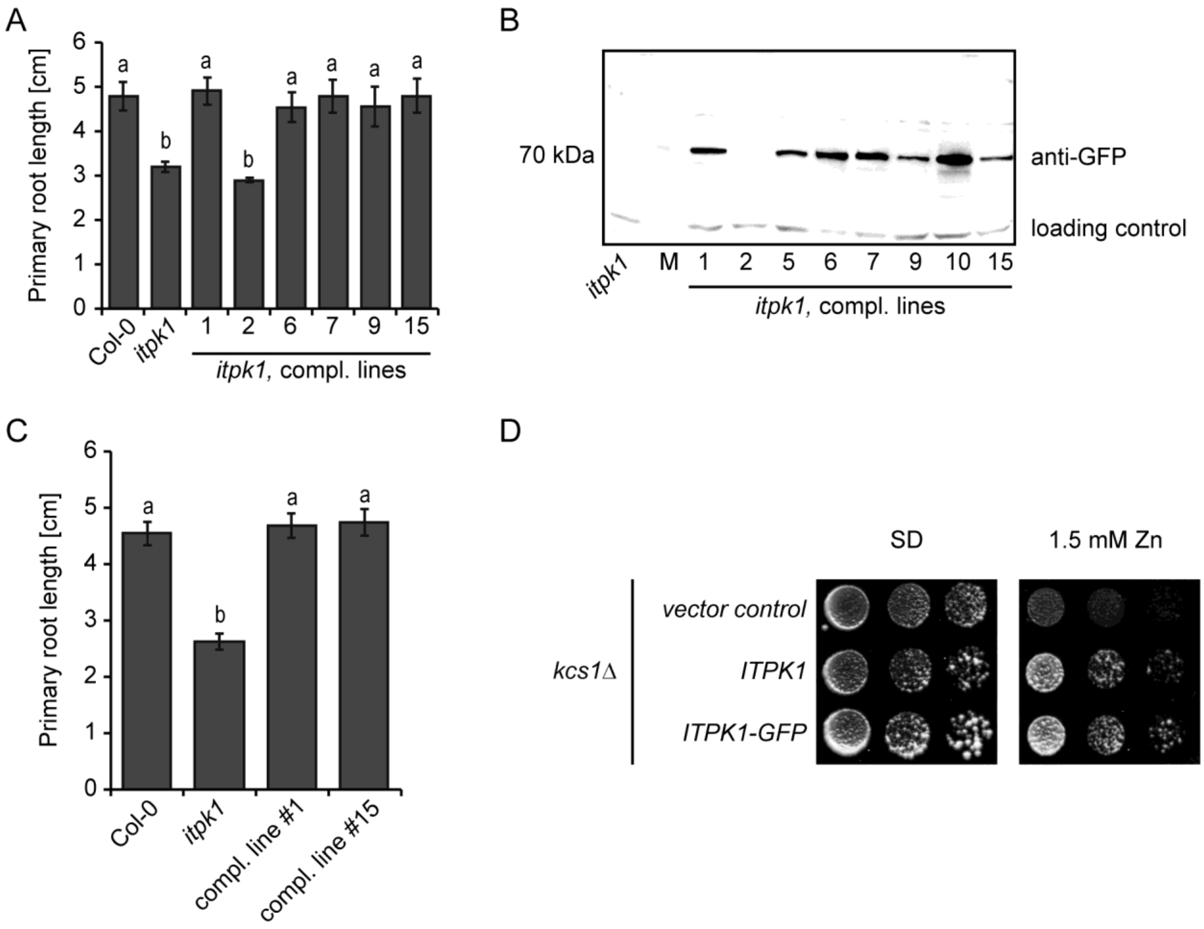


**Supplemental Figure S2.** ITPK1 regulates seedling growth and development in Arabidopsis and a C-terminal G3GFP fusion of ITPK1 does not compromise ITPK1 functions.

**(A)** Evaluation of primary root growth in wild-type (Col-0), *itpk1* mutant and independent *itpk1* lines complemented with a genomic ITPK1 fragment C-terminally fused to G3GFP. Seeds of indicated genotypes were surface sterilized and sown on sterile solid 0.5 x MS media supplemented with 1 % sucrose. Germinated seedlings were allowed to grow for 13 days and digitally recorded afterwards. Root lengths were evaluated by ImageJ. Error bars represent standard errors (s.e.), n ≥ 7. Letters indicate significance in one-way ANOVA) (a and b, $P < 0.005$). The experiment was performed twice with similar results.
**(B)** Screening for *itpk1* complemented lines. Immunoblot analyses of soluble protein extracts from 2-week-old seedlings of the indicated genotype. An unspecific band was selected as a loading control.
**(C)** Complementation of short primary root elongation of *itpk1* plants by an *ITPK1* genomic fragment. 6-day-old seedlings of designated genotypes were transferred to 0.5 x MS supplemented with 1 % sucrose and allowed to grow for another 10 days. Error bars represent standard errors (s.e.), n≥ 23. Letters depict significance in a one-way ANOVA (a and b, $P < 0.001$). The experiment was repeated twice with similar results.
**(D)** A C-terminal G3GFP fusion of ITPK1 does not compromise ITPK1 functions. ITPK1 or ITPK1-G3GFP were expressed from episomal plasmid pDR195 in a *kcs1Δ* yeast strain.

Transformants were spotted either on selective minimal medium with appropriate supplements (SD, left), or SD medium containing 1.5 mM $ZnSO_4$ (right).

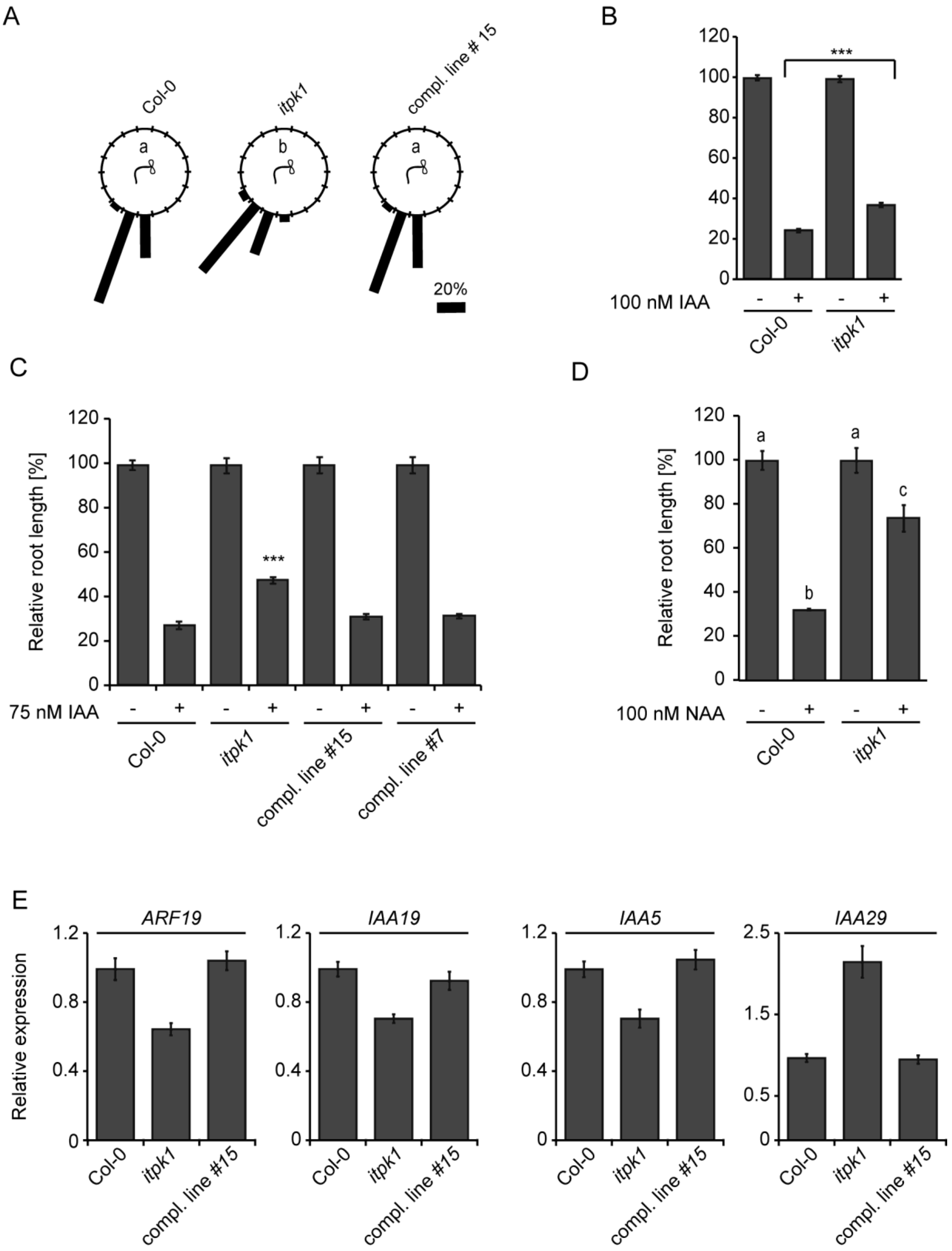


**Supplemental Figure S3.** Plant responses to exogenous auxin are regulated by ITPK1.

**(A)** Root gravitropism of seedlings of wild-type (Col-0), *itpk1* mutant and a selected complemented *itpk1* line after 90° reorientation. 7-day-old seedlings were transferred to solid plant media and after another 7 days of growth, the seedlings of indicated genotypes were rotated by 90° and the gravitropic curvature was measured after 28 h. The distribution of data was analyzed using a $\chi 2$ test (number of seedlings $n \geq 26$, groups contained at least 3.3% of total seedlings per genotype). Means with different letters are significantly different, $P < 0.001$. This represents an independent set of experiments from the experiment presented in Figure 6B. Genotypes in all panels are as indicated.
**(B)** Relative root length of wild-type (Col-0) and *itpk1* mutant treated with 100 nM IAA. Seeds of indicated genotypes were surface sterilized and sown on sterile solid 0.5 x MS, 1 % sucrose media. 6-day-old seedlings were transferred to new plates containing either 100 nM IAA or DMSO as control and scanned after 7 days. Root lengths were evaluated by ImageJ. Significant differences at $P < 0.0001$ as determined by Student's *t*-test are indicated by an asterisk. Data are shown as means ± s.e.m, $n \geq 35$.
**(C)** Relative root length of wild-type (Col-0), *itpk1* mutant and selected complemented lines treated with 75 nM IAA. 6-day-old seedlings of designated genotypes were transferred to solid 0.5 x MS, 1 % sucrose media supplemented with 0.75 nM IAA. After 7-days of growth, images were taken and root lengths were evaluated by ImageJ. Error bars present s.e.m, n=10-35. Asterisks indicate statistical differences (Student's *t*-test; ***$P < 0.0001$).
**(D)** Relative root length of wild-type (Col-0) and *itpk1* mutant treated with 100 nM NAA. Seeds of indicated genotypes were surface sterilized and sown on sterile solid 0.5 x MS, 1 % sucrose media with or without NAA. Germinated seedlings were allowed to grow for 16 days. Root lengths were evaluated by ImageJ. Letters depict significance in a one-way ANOVA (a and b, $P < 0.001$; b to c, $P < 0.001$; a to c, $P < 0.01$). Data are shown as means ± s.e.m, n=10-29.
**(E)** Relative expression of auxin-responsive genes in wild-type (Col-0), *itpk1* mutant and a selected complemented line, determined by qPCR analyses using RNA extracted from 2-week-old seedlings grown on sterile MS media. *PP2AA3* served as a reference gene. Shown are means, ± s.e.m, n=3.

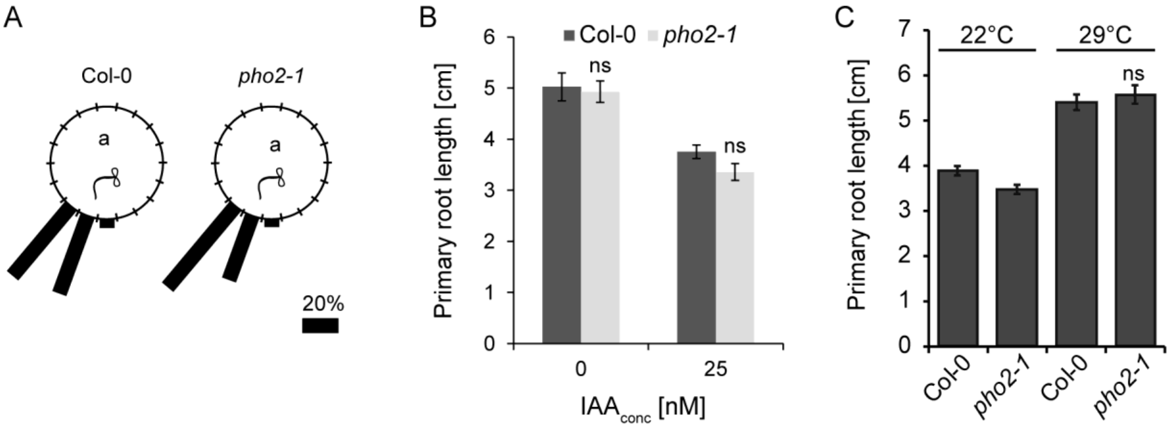


**Supplemental Figure S4.** Auxin-related growth and developmental processes are not affected in Arabidopsis *pho2-1* plants.

**(A)** Root gravitropism of seedlings of wild-type (Col-0) and *pho2-1* mutant after 90° reorientation. 7-day-old seedlings of Col-0 and *pho2-1* were transferred to solid plant media and after another 12 days of growth, the seedlings were rotated by 90° and the gravitropic curvature was measured after 16 h. The distribution of data was analyzed using a $\chi 2$ test (number of seedlings $n \geq 22$, groups contained at least 4% of total seedlings per genotype). Same letter (a) denotes that no significant differences at $P < 0.05$ were detected.
**(B)** Effect of auxin on the primary root length of the phosphorus overaccumulator mutant *pho2-1*. 6-day-old seedlings of wild-type (Col-0) and *pho2-1* were transferred to 1/2 MS plates supplemented with 0 or 25 nM IAA and 625 µM phosphate. Primary root length was measured 7 days after transferring plants to treatments. Bars show means ± s.e.m (n = 11) . No significant differences at $P < 0.05$ were detected by Student's *t*-test.
**(C)** 5-day-old seedlings of wid-type (Col-0) and *pho2-1* grown at 22°C were kept at 22°C or shifted to 29°C. Root length was evaluated after 8 days by ImageJ. Error bars represent s.e.m, $n \geq 23$. No significant differences at $P < 0.05$ were detected by Student's *t*-test.

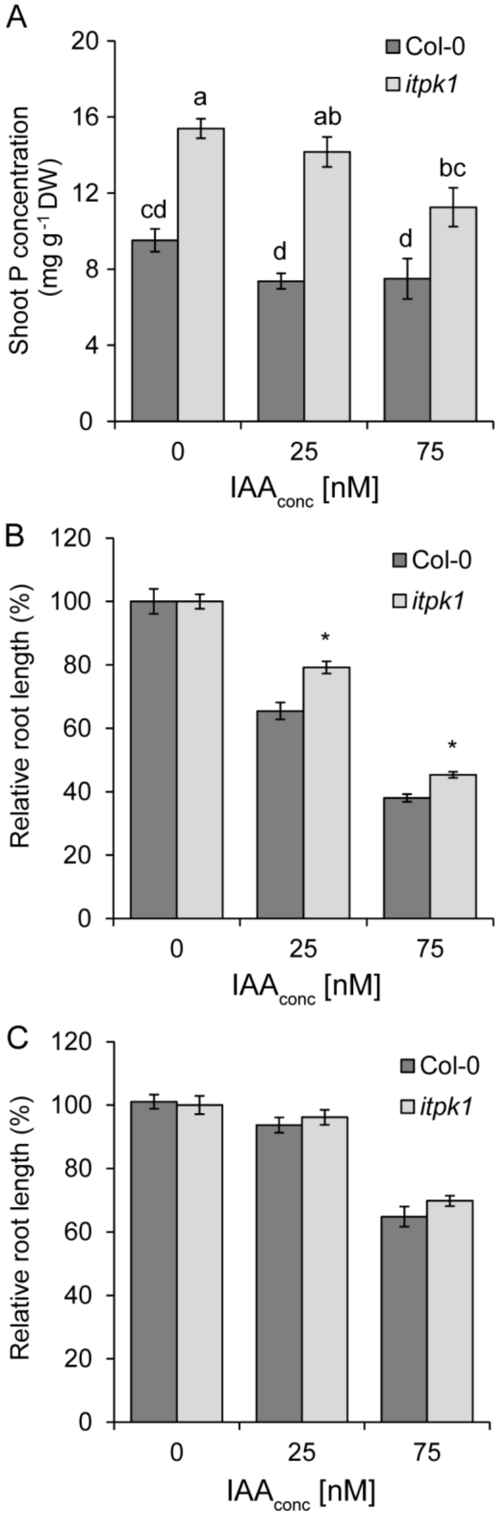


**Supplemental Figure S5.** Role of ITPK1 in phosphorus accumulation and the effect of auxin.

**(A-C)** Shoot phosphorous (P) concentration and relative primary root lengths of wild-type (Col-0) and *itpk1*mutant grown under increasing IAA concentrations. 6-day-old seedlings were transferred to 1/2 MS plates supplemented with 0, 25 and 75 nM IAA under 625 µM phosphate (*A* and *B*) or 10 µM phosphate (C). Shoot phosphorus concentration (A) and relative primary root length (B and C) were assessed 7 days after transferring plants to treatments. Bars show means ± s.e.m (n = 4 replicates with 4 plants each for shoot phosphorus analysis and 15 individual plants for primary root length). Absolute values for phosphorus concentrations were compared by ANOVA and post-hoc Tukey test and different letters indicate significant differences at $P < 0.05$. Relative root growth was compared by pairwise Student's *t*-test and significant differences at $P < 0.01$ are indicated by three asterisks.

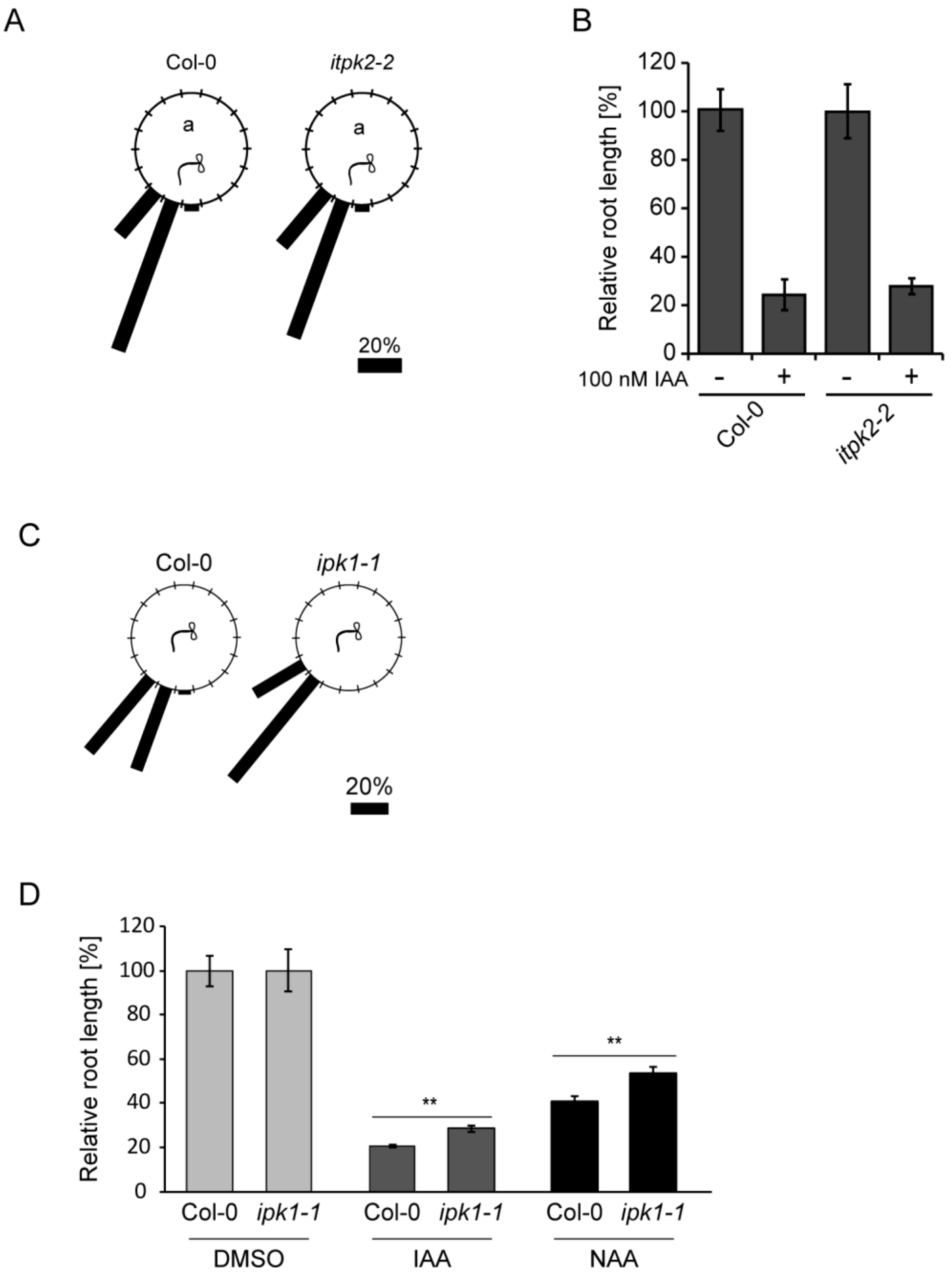


**Supplemental Figure S6.** The *ipk1-1* but not *itpk2-2* mutant is defective in auxin perception.

**(A)** Root gravitropism of seedlings of wild-type (Col-0) and *itpk2-2* plants after 90° reorientation. 7-day-old seedlings were transferred to solid plant media and after another 12 days of growth, the seedlings of Col-0 and *itpk2-2* were rotated by 90° and the gravitropic curvature was measured after 16 h. The distribution of data was analyzed using a $\chi 2$ test (number of seedlings $n \geq 35$). No significant differences at $P < 0.05$ were detected.
**(B)** Relative root length of wild-type (Col-0) and *itpk2-2* mutant treated with 100 nM IAA. Seeds of indicated genotypes were surface sterilized and sown on sterile solid 0.5 x MS, 1 % sucrose media. 6-day-old seedlings were transferred to new plates containing either 100 nM IAA or

DMSO as control and scanned after 7 days. Root lengths were evaluated by ImageJ. Data are means ± s.e.m, n=37. The experiment was repeated independently with similar results. **(C)** Root gravitropism of seedlings of wild-type (Col-0) and *ipk1-1* mutant after 90° reorientation. 12-day-old seedlings of indicated genotypes were rotated by 90° and the gravitropic curvature was measured after 16 h. The percentage of the seedlings in each category is represented by the length of the bar. The distribution of data was analyzed using a $\chi 2$ test (number of seedlings n ≥ 20). Means with different letters are significantly different, $P < 0.005$. The experiment was done independently with similar results. **(D)** Relative root length of wild-type (Col-0) and *ipk1-1* mutant treated with IAA and NAA . Seeds of Col-0 and *ipk1-1* (Laha et al., 2015) were surface sterilized and sown on sterile solid 0.5 x MS, 1 % sucrose media with or without auxin. Germinated seedlings were allowed to grow for 9 days. Root lengths were evaluated by ImageJ. Error bars present s.e.m, n≥10. Asterisks indicate statistical differences (Student's *t*-test; ***$P < 0.001$).

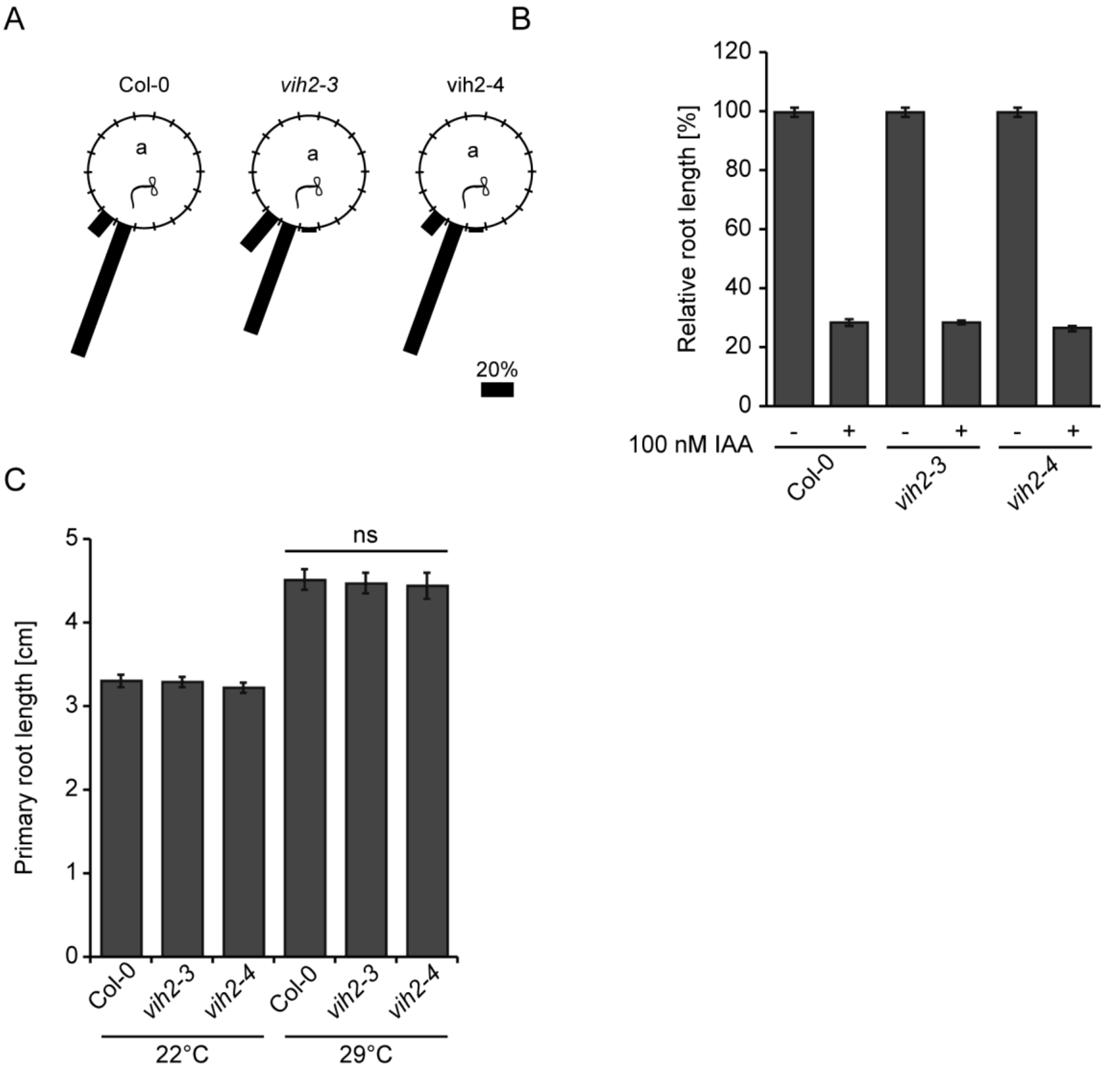


**Supplemental Figure S7.** VIH2-deficient plants are not compromised in auxin perception.

**(A)** Root gravitropism of seedlings of wild-type (Col-0), *vih2-3* and *vih2-4* mutants after 90° reorientation. 7-day-old seedlings of indicated genotypes were transferred to solid plant media and after another 12 days of growth, the seedlings were rotated by 90° and the gravitropic curvature was measured after 16 h. The distribution of data was analyzed using a $\chi 2$ test (number of seedlings $n \geq 35$). Same letter denotes that no significant differences at $P < 0.05$ were detected. The experiment was repeated independently with similar results.
**(B)** Relative root length of wild-type (Col-0), *vih2-3* and *vih2-4* mutants treated with 100 nM IAA. Seeds were surface sterilized and sown on sterile solid 0.5 x MS media supplemented with 1 % sucrose. 6-day-old seedlings were transferred to new media containing either 100 nM IAA or DMSO as control and scanned after 7 days. Root lengths were evaluated by ImageJ. Data are

means ± s.e.m, n≥34. No significant differences at $P < 0.05$ were detected by Student's *t*-test. The experiment was repeated independently with similar results.

**(C)** Primary root length analysis of seedlings of wild-type (Col-0), *vih2-3* and *vih2-4* mutants grown at higher temperatures. 5-day-old seedlings of designated genotypes were grown at 22°C, then kept at 22°C or shifted to 29°C. Root length was evaluated after 8 days by ImageJ. Error bars represent s.e.m, n ≥ 25. No significant differences at $P < 0.05$ were detected by Student's *t*-test. The experiment was repeated independently with similar results.

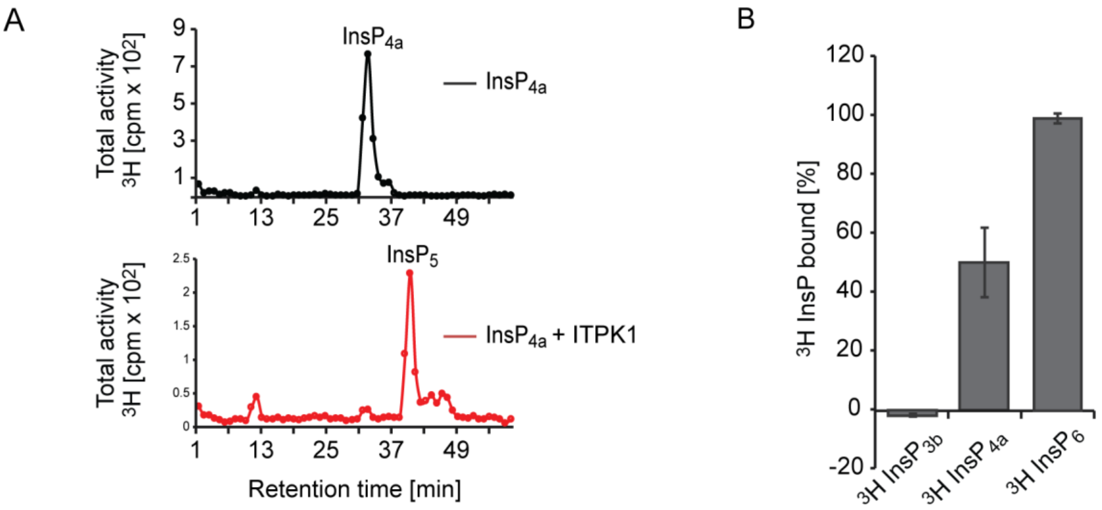


**Supplemental Figure S8.** Binding of inositol polyphosphates to the auxin-receptor complex.

**(A)** SAX-HPLC of ITPK1 kinase reaction. [$^3$H]-InsP4a was purified from [$^3$H] inositol-labeled *itpk1-2* plants and incubated with recombinant ITPK1 and ATP. The kinase product was resolved by SAX-HPLC.

**(B)** Direct binding of [$^3$H]-$InsP_{3b}$, [$^3$H]-$InsP_{4a}$, and [$^3$H]-$InsP_6$ to the TIR1/ASK1/IAA7 auxin receptor. A total activity of 2000 cpm was used for each [$^3$H]-labeled inositol phosphate species. [$^3$H]-$InsP_{3b}$ and [$^3$H]-$InsP_{4a}$ were purified and desalted from [$^3$H]-*myo*-inositol labeled seedlings of the *itpk1-2* mutant and [$^3$H]-$InsP_6$ from Col-0 seedlings. Values show means ± s.e.m. (n = 2).

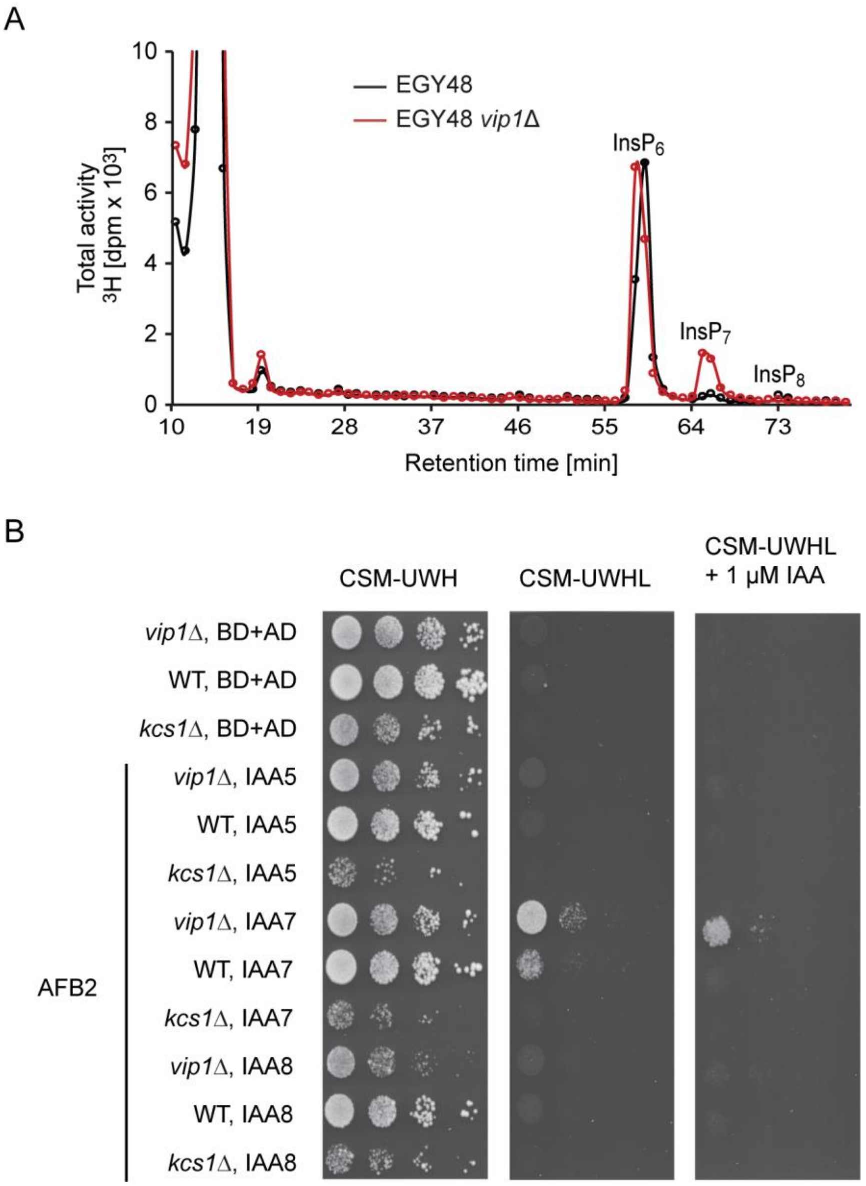


**Supplemental Figure S9.** 5-$InsP_7$ potentiates formation of auxin receptor complex *in vivo*.

**(A)** SAX-HPLC profiles of extracts of 3-week old [$^3$H] inositol-labeled yeasts. A zoom-in into the $InsP_6$-$InsP_8$ region is presented in Fig. 5A.

**(B)** Growth of yeast transformants on selective media reports interaction between AFB2 and Aux/IAA repressors. Compared to wild-type yeast transformants, *vip1Δ* transformants expressing *AFB2* and *IAA7* showed increased growth.

A

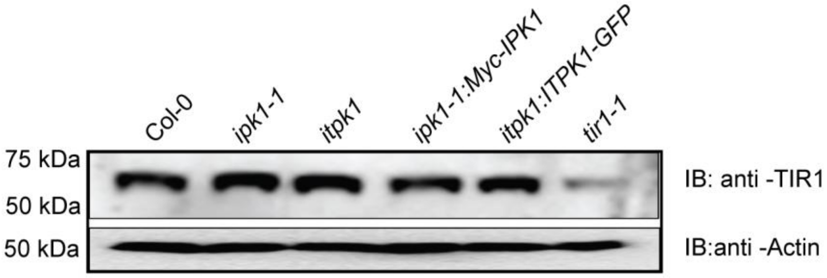


B

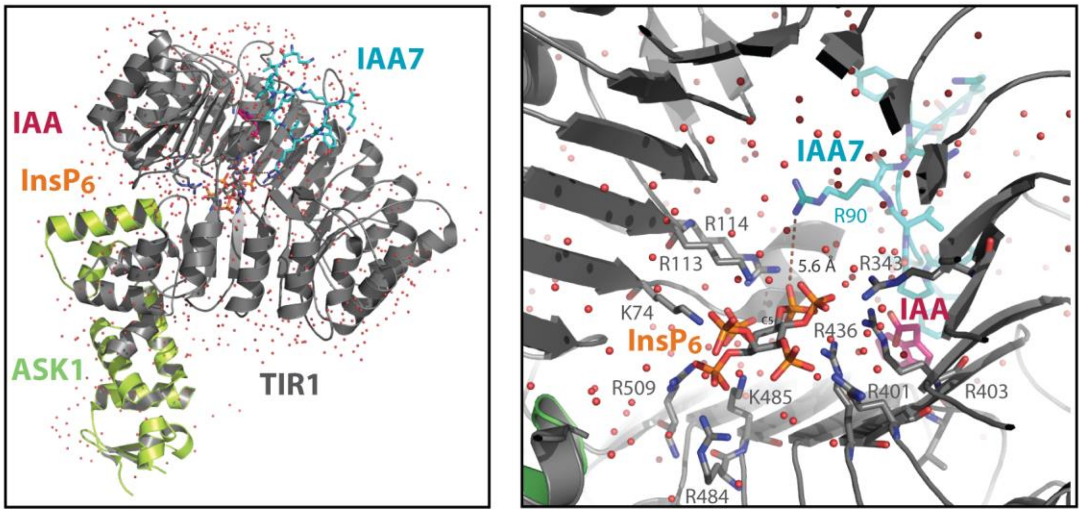


**Supplemental Figure S10.** Stability of TIR1 in *itpk1* Plants and Structural Considerations of Inositol Polyphosphate Binding of the Auxin Receptor Complex

**(A)** Levels of the auxin receptor TIR1 remain unaffected in *ipk1-1* and *itpk1* plants. Immunoblot analysis of TIR1 in whole plant extracts of 14-day-old seedlings of wild-type (Col-0), *ipk1-1*, *itpk1-2*, the respective complemented lines *ipk1-1: Myc-IPK1*, *itpk1-2: ITPK1-GFP*, and the *tir1-1* mutant. The bottom panel shows immunoblot of same extracts with anti-actin antibodies as a loading control. Migration position of molecular weight standards (in kDa) is shown at the left of each panel.

**(B)** Structural considerations of $InsP_6$ binding by the auxin-receptor complex.

Richardson diagram of the TIR1-ASK1-IAA7 degron complex bound to indole acetic acid (IAA) and $InsP_6$ (PDB ID: 2P1Q). TIR1 (gray), ASK1 (lime green), the IAA7 degron (cyan stick),

$InsP_6$ (orange stick), and IAA (magenta stick) are presented. TIR1 residues engaging in polar contacts with $InsP_6$ are depicted as sticks. Note their anisotropic distribution distal and proximal to the hormone binding pocket. The distance between IAA7 degron residue Arg 90 and the closest phosphate of $InsP_6$ (i.e. at position C5) is indicated and suggests a strong interaction with the inositol pyrophosphate moiety of 5-$InsP_7$ - provided that both $InsP_6$ and $InsP_7$ occupy the InsP binding pocket of the auxin receptor complex in a similar fashion. Images were generated with PyMOL (The PyMOL Molecular Graphics System, Version 0.99 Schrödinger, LLC).

### Supplemental Tables

#### **Supplemental Table 1.** Primer list.

Primer list for PCR-based characterization of T-DNA insertion lines.

| Mutant lines | Sense Primer (5'-3') | Anti-sense Primer (5'-3') |
|---|---|---|
| *itpk1* | GCTTCCTATTATATCTTCCCAAATTACCA ATACA (LB2_SAIL) | CATGGCTTTGCAAAACTCG GAAG |
| *itpk2-2* | GCTTCCTATTATATCTTCCCAAATTACCA ATACA (LB2_SAIL) | TCGGTTATGTTTAAACGCC AAC |

Primers used for qPCR-analyses.

| Gene | Sense Primer (5'-3') | Anti-Sense Primer (5'-3') |
|---|---|---|
| *IAA29* | GGGTGCTGCGTCTTGTTTGGGT | TCCTCGTTGGGCTGGCCATT |
| *IAA5* | GTCGTCTCCGGTGAGTCCATCT | AAACCGGTGGCCAACCCACAA |
| *PP2AA3* | GGCAGAAGTTCGGATAGCAG | CAATGCAGATCTGACGTGCT |
| *IAA19* | TGGCATCGGTGTGGCCTTGAA | ACATCACCGGCGAGCATCCAGT |
| *ARF19* | TTGAGCGCGCAAGCAATCCG | TGCCTTGACTGATCCGGCTCTCA |
| *ITPK1* | AGTTGGATACGCACTCGCAGCC | TGCCTCGTTGCCTTGAGTGTTCG |
| *ITPK2* | AACGCAGACTTGGACCCTCGTG | AAGCGCCTCGAGGAAAGGCT |

Primer list to clone into the pENTR-D-TOPO vector.

| Gene | Sense Primer (5'-3') | Anti-Sense Primer (5'-3') |
|---|---|---|
| *ITPK1* | caccATGTCAGATTCAATCCAGGAAAG | TCAGACATGATTCTTCTTAGTG |
| *ITPK2* | caccATGTTTGGGACTCTTGCTTCCGGC | TCATTTACAATGTTGTCTCTT |
| *ITPK3* | caccATGTACTGGCAGCAAATTGGA | TTAGTATTGATCTGCCAAAGCT |
| *ITPK4* | caccATGAAAGGGGTTCTACTTGACGA | TCAATGCTTCTCTTGGACATGTT |
| *IPK2a* | caccATGCAGCTCAAAGTCCCTGAAC | CTAAGAATCTGCAGACTCATC |
| *IPK2b* | caccATGCTCAAGGTCCCTGAACA | CTAGCGCCCGTTCTCAAGTAGG |
| *IPK1* | caccATGGAGATGATTTTGGAGGAGA | ttgcggccgcTTAGCTGTGGGAAGGTTTTGA G |

Primers employed for site-directed mutagenesis.

| Gene | Mutation | Primer sequence (5'-3', only sense orientation is listed) |
|---|---|---|
| *ITPK1* | K188A | CATGGTGGTGTGATCTTTgcGGTCTATGTGGTTGGAGATC |
| *ITPK1* | D288A | GCTAATAGGTACCTTATAATTGcTATTAACTACTTTCCTGG |
| *ITPK2* | K260A | AATCATGGTGGAGTTATGTTCgcGGTATTTGTGGTGGGTGATGTTA |
| *ITPK2* | D355A | GCAAAAACGTGTTTTATGTTATTGcCATCAACTATTTTCCTGGT |

Specific mutations are in lower cased. Corresponding mutations in the final plasmids were confirmed by sequencing.

Primers and plasmids used to generate yeast deletion strains.

| Gene | Primer Sequence (5'-3') | Plasmid |
|---|---|---|
| *VIP1* | AATTCAAAAGCATCTCGTAGCATATTAATATATTGCAGAAGGTCcagc tgaagcttcgtacgc (sense)<br>ACTTATTTAGTTTTGGGTTACTAAATTAAAAATTGGGTGTGATCAgcat aggccactagtggatctg (antisense) | pUG6 (Euroscarf) |

Letters in lower case indicate the primer region that recognizes the plasmid sequence.

Primers employed to clone into the pET28- $His_8$-MBP bacterial expression vector

| Gene | Primer Sequence (5'-3') |
|---|---|
| *ITPK1* | AAggatccATGTCAGATTCAATCCAGGAAAG (sense) |

| | |
|---|---|
| | TgtcgacTCAGACATGATTCTTCTTAGTG (antisense) |
| *ITPK2* | AAgaattcATGTTTGGGACTCTTGCTTCCGG (sense)<br>TgtcgacTCATTTACAATGTTGTCTCTTCT (antisense) |
| *hITPK1* | AggatccATGCAGACCTTTCTGAAAGGGAA (sense)<br>tGCGGCCGCCTACTGGGAGGAGGCCTTGGTGG (antisense) |
| *IAA7* | AggatccATGATCGGCCAACTTATGAACCT (sense)<br>TgcggccgcTCAAGATCTGTTCTTGCAGTAC (antisense) |
| *VIH2* | AAggatccATGGAGATGGAAGAAGGAGCA (sense)<br>TTgcggccgcTTAGCTCCTTCCATTAGAAGAAG (antisense) |

Primers for amplifying ITPK1 and IPK1 cDNAs

| cDNA | AttB1 For | Attb1 Rev |
|---|---|---|
| ITPK1 | 5'-GGGGACAAGTTTGTACAAAAAAGCAGGCTCAATGTCAGATTCAATTC-3' | 5'-GGGGACCACTTTGTACAAGAAAGCTGGGTATCAGACATGATTCTTCTT-3' |
| TIR1 | 5'-GGGGACAAGTTTGTACA AAAAAGCAGGCTCAATGCAGAAGCGAATAGCCTTGTCG-3' | 5'-GGGGACCACTTTGTACA AGAAAGCTGGGTATTATAATCCGTTAGTAGTAATGAT |
